# The ontogeny of myeloid-stromal synovial tissue niches in rheumatoid arthritis

**DOI:** 10.1101/2025.10.29.685324

**Authors:** Aziza Elmesmari, Domenico Somma, Lucy MacDonald, Jack Frew, Zuzanna Kuczynska, Clara Di Mario, Lavinia Agra Coletto, Shravani Baruah, Denise Campobasso, Theodoros Simakou, Eva M. L. Philippon, Sander W. Tas, Marcus Doohan, Roberta Benvenuto, Dario Bruno, Maria Rita Gigante, Luca Petricca, Viviana Antonella Pacucci, Kevin Wei, Charles P. McSharry, Dylan Windell, Sam Pledger, Sarah Davidson, Mark C Coles, Marco Gessi, David Tough, Maria Antonietta D’Agostino, Iain B. McInnes, Stephanie G. Dakin, Christopher D. Buckley, Stefano Alivernini, Mariola Kurowska-Stolarska

**Author notes:** **Corresponding authors:** Mariola Kurowska-Stolarska,; Stefano Alivernini. Equally contributing first (Aziza Elmesmari, Domenico Somma, Lucy MacDonald). last authors (Stefano Alivernini, Mariola Kurowska-Stolarska).

## Abstract

Recent single-cell multi-omic and spatial analyses of synovial biopsies have transformed our understanding of myeloid cell–driven mechanisms underlying human joint pathology and tissue homeostasis in Rheumatoid arthritis (RA). However, the developmental trajectories of synovial tissue macrophage (STM) subsets in humans remain poorly understood, due in part to the lack of models that faithfully replicate synovial tissue niches. This hinders the exploration of the therapeutic potential of targeting specific synovial myeloid cell clusters. Using multi-omics analyses of synovial tissue from an allogeneic bone marrow transplant recipient, we show that joint-specific tissue-resident STM subsets, including both health- and disease-associated clusters, can derive from peripheral blood monocytes. Analysis of embryonic synovial joints revealed that macrophage localization and maturation in the joints are preceded by local stromal niche specialisation, indicating that synovial fibroblasts (FLS) provide tissue-specific instructive cues to STM precursors. To elucidate human STM developmental trajectories, we established a SNP-based fate-tracking human synovial organoid system by embedding distinct blood-derived myeloid precursors, together with FLS clusters from RA synovial biopsies and endothelial cells, into 3D structures. These organoids reproduced key synovial tissue features, including lining and sublining architecture and stromal–myeloid cell cluster composition. Importantly, they supported differentiation of all resident STM subsets: homeostatic lining TREM2^pos^ macrophages, their pathogenic TREM2^low^SPP1^pos^ counterparts that characterize the RA hyperplastic lining, and both homeostatic and RA-associated perivascular LYVE1^pos^ STM clusters, all traced to monocytic precursors. In summary, we show that development of STM subsets is driven by fibroblast-conditioned spatial niches. We have established a novel, tractable ex vivo platform to dissect the niche-specific cues driving homeostatic versus pathogenic phenotypic clusters.

**One Sentence Summary:** Human tissue-resident STMs develop from monocytes under the guidance of cues from FLS within discrete spatial locations.

## INTRODUCTION

High-resolution omics tissue biology, combined with *ex vivo* functional studies and animal models, have identified a diversity of macrophage subsets in human synovial tissue with distinct roles in joint homeostasis and inflammation, transforming understanding of Rheumatoid arthritis (RA) pathology and remission (*1*). However, the origins and mechanisms underlying the development of these populations in the human synovium in health and RA remain unclear, limiting their potential for specific therapeutic targeting.

The synovial membrane forming the inner layer of the joint capsule, is a specialized tissue composed of two distinct layers: a lining layer (LL) that faces the joint cavity and a sublining layer (SL) containing sensory neurons and blood vessels (*1, 2*). Together, these layers provide immune-protection and joint lubrication. Within all organs, reciprocal communication between tissue macrophages and fibroblasts is crucial, with the outcome of these interactions often determining organ function, including that of synovial joints (*3*). For clarity, throughout this study we refer to synovial macrophage subsets that are present in healthy joints and persist across various conditions as tissue-resident subsets (populations). These subsets can adopt different phenotypic states (clusters), displaying homeostatic/regulatory functions in health or during sustained disease remission, or acquiring pathogenic states in active disease. In contrast, macrophage populations/clusters that appear only in inflamed synovium and resolve during disease remission are classified as non-resident (*4*).

In healthy joints, tissue-resident lining MerTK^pos^TREM2^pos^VSIG4^pos^ synovial tissue macrophage population (STMs) form an epithelial-like barrier structure (*5*), tightly interacting with lining-layer PRG4^pos^PDPN^pos^CLU^pos^CD90^neg^ synovial fibroblasts (FLS) (*6*) and tolerogenic myeloid AXL^pos^ dendritic cells (DC2) to support lubrication, control adaptive immunity and resolve pathology (*5, 7, 8*). Both lining layer STMs (*2*) and FLS (*9*) also communicate with axons of sensory neurons that extend into the lining niche, mediating pain when joint homeostasis is lost (*2*). In the sublining, tissue-resident MerTK^pos^ perivascular LYVE1^pos^ STM populations, together with perivascular CD90^pos^CD34^pos^ and CD90^pos^GAS6^pos^ FLS and sensory neurons, regulate vascular permeability as well as the recruitment and activation of inflammatory cells (*2, 4, 10*). Additionally, collagen-producing synovial fibroblast clusters (e.g. POSTN^pos^COL1/3^pos^) provide the structural integrity of the tissue (*6, 11*). In rheumatoid arthritis, the synovial lining shows a decline in protective TREM2^pos^ STMs, loses its homeostatic function, and becomes hyperplastic (*7*). Lining fibroblasts adopt a destructive phenotype, producing cartilage- and bone-degrading enzymes (*6, 7*). Tolerogenic AXL^pos^ DC2 are replaced by FABP5^pos^ inflammatory DC3 (iDC3), creating a niche that promotes local T-cell activation (*12*). In parallel, in the sublining, perivascular LYVE1^pos^ STMs undergo phenotypic changes, producing chemokines themselves (*13*) or inducing neighbouring perivascular FLS to secrete cytokines and chemokines (*7*), thereby recruiting inflammatory cells. This is associated with the emergence of pathogenic non-resident MerTK^neg^ monocyte/macrophage subsets, including S100A12^pos^, ISG15^pos^, ICAM1^pos^TNF^pos^, and FCN1^pos^SPP1^pos^ STMs, which secrete TNF together with cluster-specific mediators (alarmins, IFNs, chemokines, and SPP1), suggesting distinct pathogenic functions within discrete disease associated tissue niches. Collectively, these cells amplify inflammation and drive the expansion of sublining CD90^pos^ fibroblast clusters, thereby promoting arthritis chronicity (*6, 7*).

We recently showed that RA patients who achieve sustained disease remission after treatment withdrawal (∼20% of patients), restore functional homeostatic STM subsets and eliminate pathogenic subsets (*7*). To extend this outcome to a broader RA population, a better understanding of the origins and developmental trajectories of distinct homeostatic and pathogenic STM subsets within their tissue niches is needed. In mice, fate-mapping studies have shown that protective lining TREM2^pos^MerTK^pos^VSIG4^pos^ macrophages originate from foetal liver precursors and are maintained in adulthood through in situ self-renewal (*5*). Recent evidence suggests that, during the resolution of experimental arthritis, they can be partially reinstated from blood monocyte precursors (*13*). In humans, however, the ontogeny of tissue-resident STM clusters remains largely unknown.

Herein, we elucidated the origins and developmental trajectories of distinct human STMs by examining synovial joints from embryonic through adult stages, as well as from patients with active RA and those in remission, combining SNP-based cell fate tracking with a myeloid multi-precursor synovial organoid model.

## RESULTS

### TREM2^low^SPP1^pos^ macrophages define the hyperplastic lining niche in active RA

Prior to dissecting the developmental trajectories of human STM subsets, we first aimed to addressed key gaps in understanding the phenotypes of STMs in patients with active RA. While the barrier-like regulatory phenotype of lining layer MerTK^pos^TREM2^pos^VSIG4^pos^ macrophages in healthy joints and RA in remission is well characterized, the phenotype of macrophages in the diseased hyperplastic lining layer, which serves as the niche for the recently described iDC3-driven activation of CCL5-producing memory T-cells in RA (*12*), remains poorly understood. To address this gap, we generated an extended single-cell transcriptomic atlas of synovial tissue myeloid cells across health, active RA, and RA in sustained remission **(Fig. 1A–C).** We integrated our recent datasets (*7, 12*) with scRNAseq data from additional synovial biopsies (n = 16) and matched blood myeloid cells, expanding the synovial tissue myeloid cell dataset to 60,507 cells from 45 individuals: healthy controls (n = 11), treatment-naïve active RA (n = 12), active RA resistant to conventional (c) or biologic (b) DMARD therapy (n = 11), and RA in sustained remission (n = 11) (**Fig. 1C; figs. S1–S2).**

**Figure 1.**
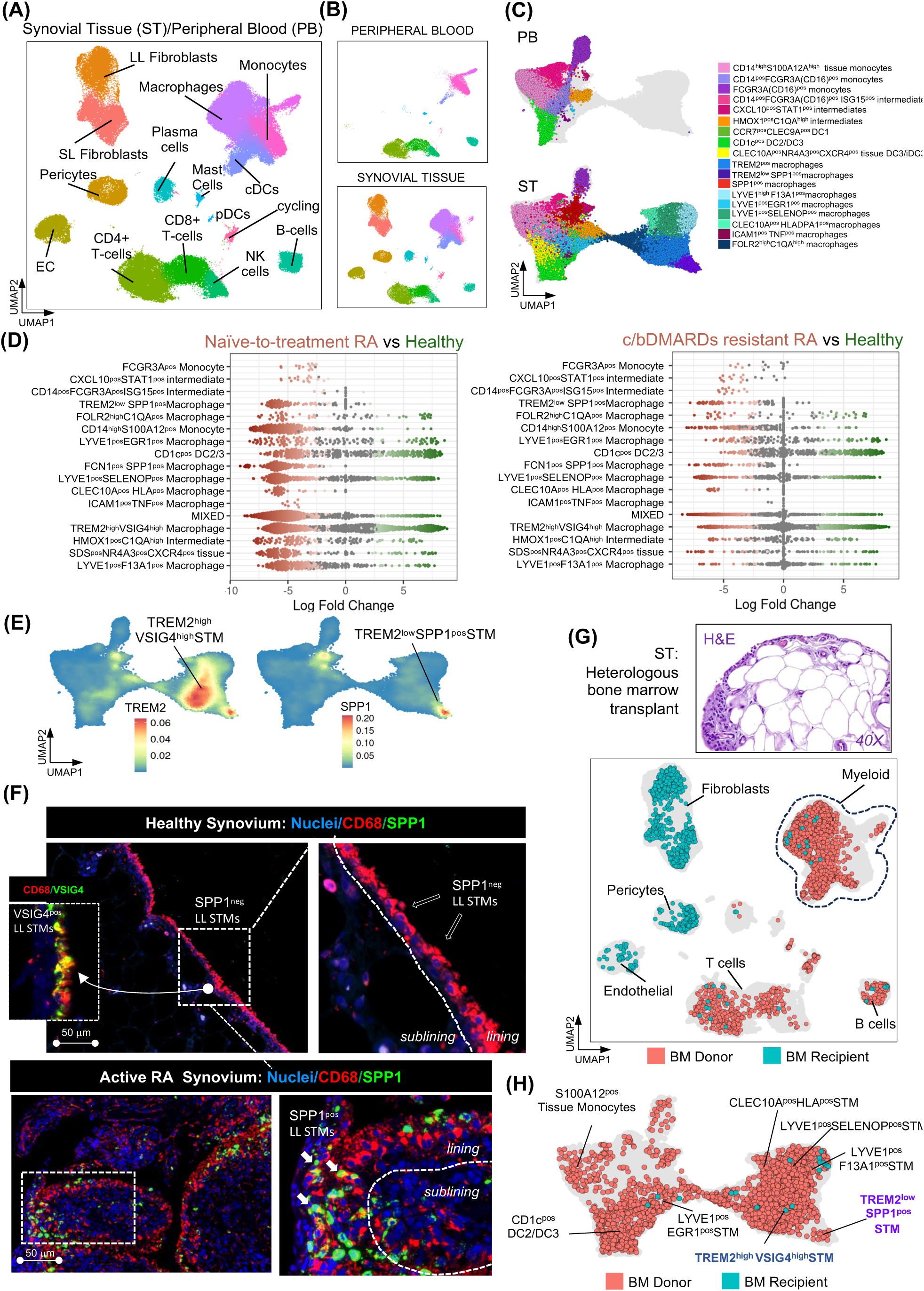
PB Monocytes can give rise to all synovial tissue–resident macrophage clusters, including lining TREM2^pos^ and its pathogenic SPP1^pos^ phenotype. **(A)** UMAP of integrated single-cell transcriptomic data from synovial tissue (ST) and matched peripheral blood (PB) of a patient with heterologous bone marrow (BM) transplant (BMT), and a reference dataset of synovial tissue from healthy controls (n=11), treatment-naïve RA (n=12), c/bDMARD-resistant RA (n=11), and RA in sustained remission (n=11). **(B)** Same as in (A) but split into PB (n= 8, 20,073 cells, and ST compartments n=45, 60,507 cells). **(C)** UMAP visualization of integrated single-cell transcriptomic data of ST myeloid cells and matched PB myeloid cells of a patient with a heterologous bone marrow transplant, and a reference ST/PB dataset from RA patients and healthy controls as in A. **(D)** Differential abundance showing changes in cell population frequencies between conditions. Data are visualized as MiloR neighbourhood graphs, where nodes represent neighbourhoods, coloured by their log fold change across conditions. Neighbourhoods with non-differential abundance (FDR > 10%) are coloured grey, and node size reflects the number of cells in each neighbourhood. **(E)** Density plots showing VSIG4 and SPP1 expression in myeloid cell atlas as in C. **(F)** Immunofluorescence images showing expression of CD68, VSIG4 and SPP1 in synovial tissue biopsies of healthy (HC) and of naïve-to-treatment RA patient, representative of at least n=3 healthy and n=3 RA patients. CD68 red; SPP1, and VSIG4 green (RNAscope); and Nuclei, blue. Scale bars, 50µm. **(G)** Histology of the synovial biopsy and UMAP visualization of synovial tissue cells from a heterologous BM transplant patient, showing donor-or recipient-derived origin as determined by SNP deconvolution (see Methods). **(H)** Visualization of synovial tissue myeloid cells from a patient with a heterologous BM transplant on the UMAP of the integrated synovial tissue myeloid cell dataset shown in (C), indicating whether the synovial myeloid cells are derived from the bone marrow donor or recipient.

Using validated synovial myeloid cell annotations (*7, 12, 14, 15*), we compared the abundances of distinct myeloid cell phenotypes between healthy and active RA joints, as well as between active RA and RA in sustained remission, using MiloR, which estimates phenotype enrichment via neighbour-based grouping (*16*). We confirmed enrichment of MerTK^neg^ non-resident clusters, including S100A12^pos^, ISG15^pos^, ICAM1^pos^TNF^pos^, and FCN1^pos^SPP1^pos^ cells, in active RA, particularly in treatment-naïve patients, with most of these resolved in sustained remission (**Fig. 1D and figs. S1–S2).** Beyond these, this expanded atlas revealed additional disease-associated phenotypes enriched in active RA, especially among tissue-resident lining TREM2^pos^, perivascular LYVE1^pos^, and antigen-presenting CLEC10A^pos^ populations. Notably, we observed a marked expansion of a phenotype within the lining TREM2^pos^ population that exhibited high expression of the pathogenic mediator SPP1 **(Fig. 1E),** which we recently identified as a potent driver of neutrophil activation and alarmin/cytokine release by monocytes (*17*).

Immunofluorescence staining of synovial biopsies from healthy controls and patients with active RA confirmed this finding at the spatial level: while healthy VSIG4^pos^ lining layer STMs were SPP1-negative, macrophages in the hyperplastic lining of active RA patients were strongly SPP1-positive, establishing this TREM2^low^ SPP1^pos^ cluster as a defining feature of the pathogenic lining niche **(Fig. 1F).**

### Bone marrow monocytes give rise to all synovial tissue–resident macrophage populations

To investigate the precursor lineage of distinct synovial tissue macrophages identified in health and across the trajectory of RA **(Fig. 1A–D)**, we performed an ultrasound-guided synovial biopsy in a patient with arthralgia (joint pain) who had undergone a heterologous bone marrow transplant (BMT) four years earlier. To facilitate cell annotation, single-cell transcriptomic profiles of the synovium and matched peripheral blood (PB) from the BMT patient were integrated with our human PB/ST single-cell reference atlas **(Fig. 1A–C**). To distinguish recipient-from donor-derived cells, we applied single-nucleotide polymorphism (SNP) deconvolution using Souporcell v2.4 (*12*). As expected, synovial tissue stromal cells, such as synovial fibroblasts, endothelial cells, and pericytes, originated from the recipient, whereas T and B cells were derived from donor bone marrow **(Fig. 1G).** Detailed SNP-based deconvolution of each synovial tissue myeloid cluster revealed that all resident populations originated from donor bone marrow. This included the lining layer TREM2^pos^VSIG4^pos^ STMs and the sublining perivascular LYVE1^pos^F13A1^pos^ populations, as well as their clusters whose activation is associated with RA, namely, the hyperplastic lining TREM2^low^SPP1^pos^ cluster and the perivascular LYVE1^pos^EGR1^pos^ and LYVE1^pos^SELENOP^pos^ clusters **(Fig. 1H).** As expected, non-resident MerTK^neg^ tissue monocyte/macrophage clusters (e.g., S100A12^pos^, ISG15^pos^, ICAM1^pos^TNF^pos^, and FCN1^pos^SPP1^pos^), albeit present in limited numbers in non-inflamed BMT synovium **(fig.S1D)**, were also of monocytic origin.

Altogether, this suggests that all synovial tissue macrophage populations in humans, including those considered to be locally self-renewing from prenatal precursors in mice (e.g., lining layer TREM2^pos^ STMs), can originate from bone marrow–derived monocytic precursors.

### Sequential embryonic development of joint-specific synovial niche

Next, to investigate how myeloid precursors acquire synovial tissue–specific phenotypes, we examined the sequence of synovial tissue niche establishment during human embryonic joint development. We recently characterized the stromal composition of early embryonic synovial joints (8–10 weeks post-conception, PCW) and identified both early and postnatal progenitors of human synovial fibroblasts (*18*). To determine when synovial tissue–resident macrophages first appear in the synovium, we stained human embryonic joints at 9–10 and 13–14 weeks post-conception for CD68 (pan-macrophage marker), TREM2 (lining macrophage marker), and PRG4 (lining fibroblast marker), as well as CD31 (blood vessel marker) to localize the sublining layer. At 9–10 PCW, the synovium was developmentally immature, lacking the characteristic organization into lining and sublining layers and mostly devoid of macrophage populations **(Fig. 2A).** However, by 13–14 PCW, a distinct lining layer of PRG4^pos^ fibroblasts was established, along with blood vessels in the sublining. While most of the synovium remained macrophage-free, specific areas of the PRG4^pos^ lining layer began to show the emergence of TREM2^pos^ macrophages at 14 PCW **(Fig. 2B).**

**Figure 2.**
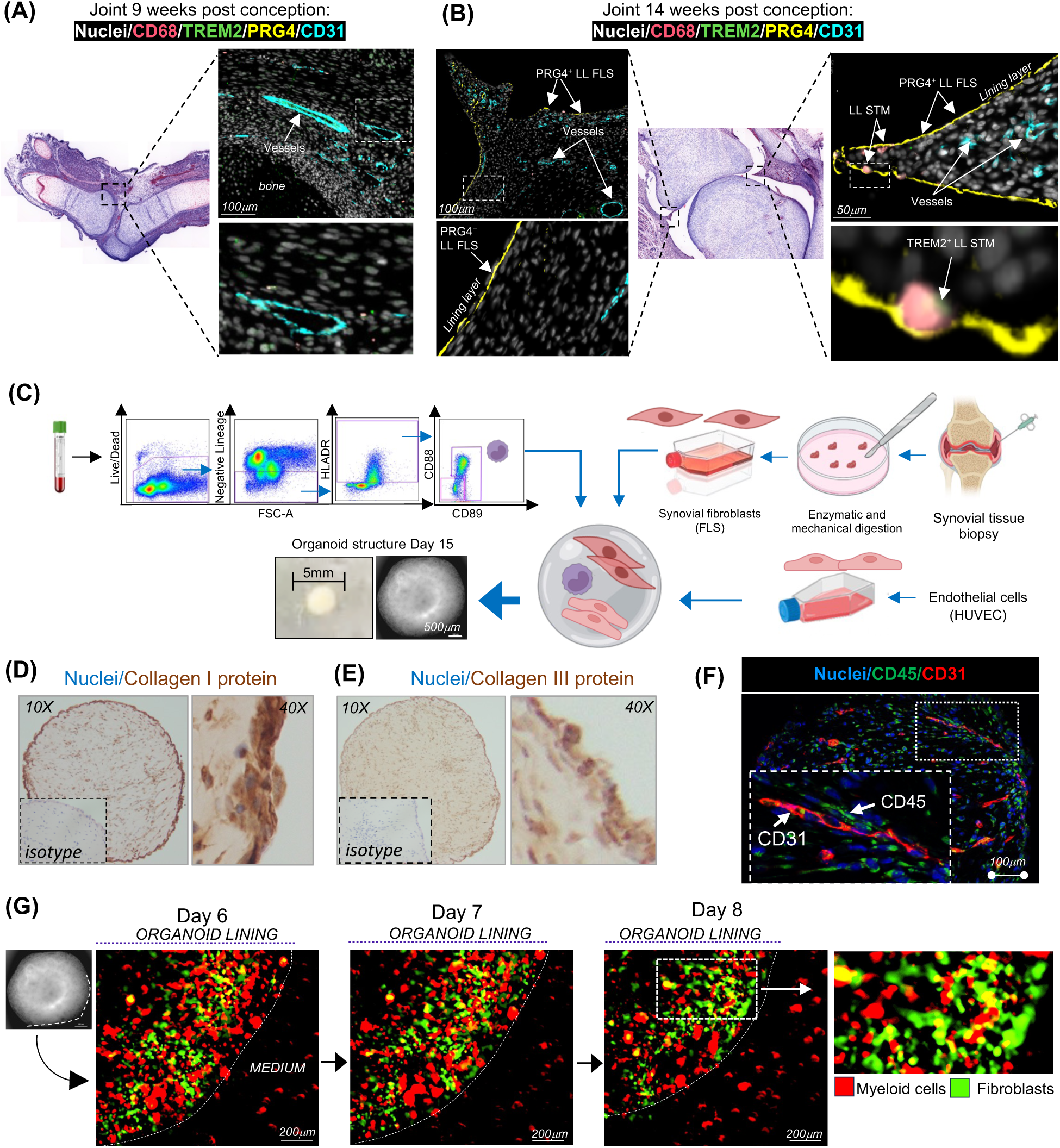
Establishment of synovial tissue niches in human embryonic joint and organoids. **(A-B)** Immunofluorescence staining of PRG4 (yellow), CD68 (red), CD31 (cyan), TREM2 (green) and nuclei (DAPI, white) in human embryonic joints at 9-14 weeks post-conception (pcw), representing data from two joints at 9-10 pcw (A) and four joints at 13-14 pcw (B). **(C)** Schematic illustrating the workflow of synovial organoid (SO) generation using synovial fibroblasts, PB monocytes, and endothelial cells in a single droplet of Matrigel. All monocyte subsets were sorted from the PB of healthy donors by excluding dead cells and lineage-positive cells, followed by gating based on HLADR and CD88/CD89 expression. Synovial fibroblasts were isolated from synovial biopsies of patients with active RA, while endothelial cells were human umbilical vein endothelial cells (HUVECs). The macroscopic appearance of the SO structure, with clearly defined edges, was observed under bright-field microscopy using a Nikon inverted microscope. **(D-E)** Representative images of IHC staining of the SO for Collagen I (D) and Collagen III (E) from organoids with FLS derived from biopsies of n=6 patients with active RA. The inserts show isotype controls. The entire SO is shown at 10x magnification, while the enlarged regions of the LL of the SO are shown at 40x magnification. Collagens are brown, and nuclei are blue. **(F)** Representative confocal microscopy images (40x) showing IF staining for CD31 (red), macrophages (CD45, green) and nuclei stained with DAPI (blue) from organoids with FLS derived from biopsies of n=6 patients with active RA at different time points. The insert shows enlarged region of SO. Scale bars, 100 µm. **(G)** Representative images of myeloid cell-FLS interactions in the synovial organoids (SO), generated using CellTracker Red-labelled PB monocytes and CellTracker Green-labelled synovial fibroblasts. These images are derived from continuous live recordings conducted using the Omni imaging system at different time points. Data representative of one experiment with three technical replicates.

These findings suggest that the joint stromal niche matures first, subsequently providing tissue-specific cues that guide myeloid precursor localization and differentiation into STMs.

### Myeloid-synovial stroma organoids recapitulate synovial tissue topography

Next, to investigate the role of synovial tissue stroma as the most likely source of tissue-specific cues for STMs, we have developed synovial organoid model incorporating myeloid cell precursors and synovial tissue biopsy–derived stroma into 3D structure. To specifically assess stromal contributions while excluding potential inflammatory imprinting of bone marrow or blood precursors, we used healthy donor myeloid precursors rather than those from RA patients. After excluding other lineage-positive cells, we sorted blood monocytes based on their unique expression of CD88/CD89 (*12, 19*) and co-embedded them together with synovial fibroblast clusters derived from synovial biopsies of active RA patients and endothelial cells (HUVEC cell line) in Matrigel **(Fig.2C)**. This led to the formation of sphere-like organoids within 15 days, with progressive growth **(fig.S3A-C)** and sustained cell viability up to 21 days of monitoring **(fig.S3D).** Staining for extracellular matrix components and endothelial cells demonstrated the stromal functionality, including the production of collagen I and III (absent in Matrigel) **(Fig.2D-E)** and the formation of tube-like endothelial structures **(Fig.2F and fig.S3E)**, collectively indicating the establishment of a viable stromal niche.

To determine whether myeloid blood precursors were able to interact with the stroma in the organoids, blood monocytes and biopsy-derived FLS were labelled with red and green cell trackers, respectively, prior to embedding in Matrigel, and monitored by live-imaging for 8 days **(Fig.2G).** This revealed that myeloid cells closely interact with synovial fibroblasts within the organoids, inferring active cell-to-cell communication. Active RA tissue is characterized by an abnormal lining, in which PRG4^pos^ FLS adopt a pathogenic, MMP3-producing phenotype **(Fig. 3A)**, and a sublining layer containing expanded CD90^pos^ (THY1) FLS and blood vessels (*6*), with macrophages present in both niches but particularly enriched in the lining **(Fig. 3A).** Staining of myeloid–synovial stromal organoids with antibodies against lining and sublining markers, together with mapping of myeloid cells using anti-CD45 antibody, showed that these organoids developed a distinct lining layer containing PRG4^pos^MMP3^pos^ FLS **(Fig. 3B–C)** and a sublining layer with CD90^pos^ FLS and blood vessel–like structures **(Fig. 3D and fig. S3E)**, with macrophages present in both niches **(Fig. 3E)**. This tissue-like topography was established by day 15 and maintained through day 29.

**Figure 3.**
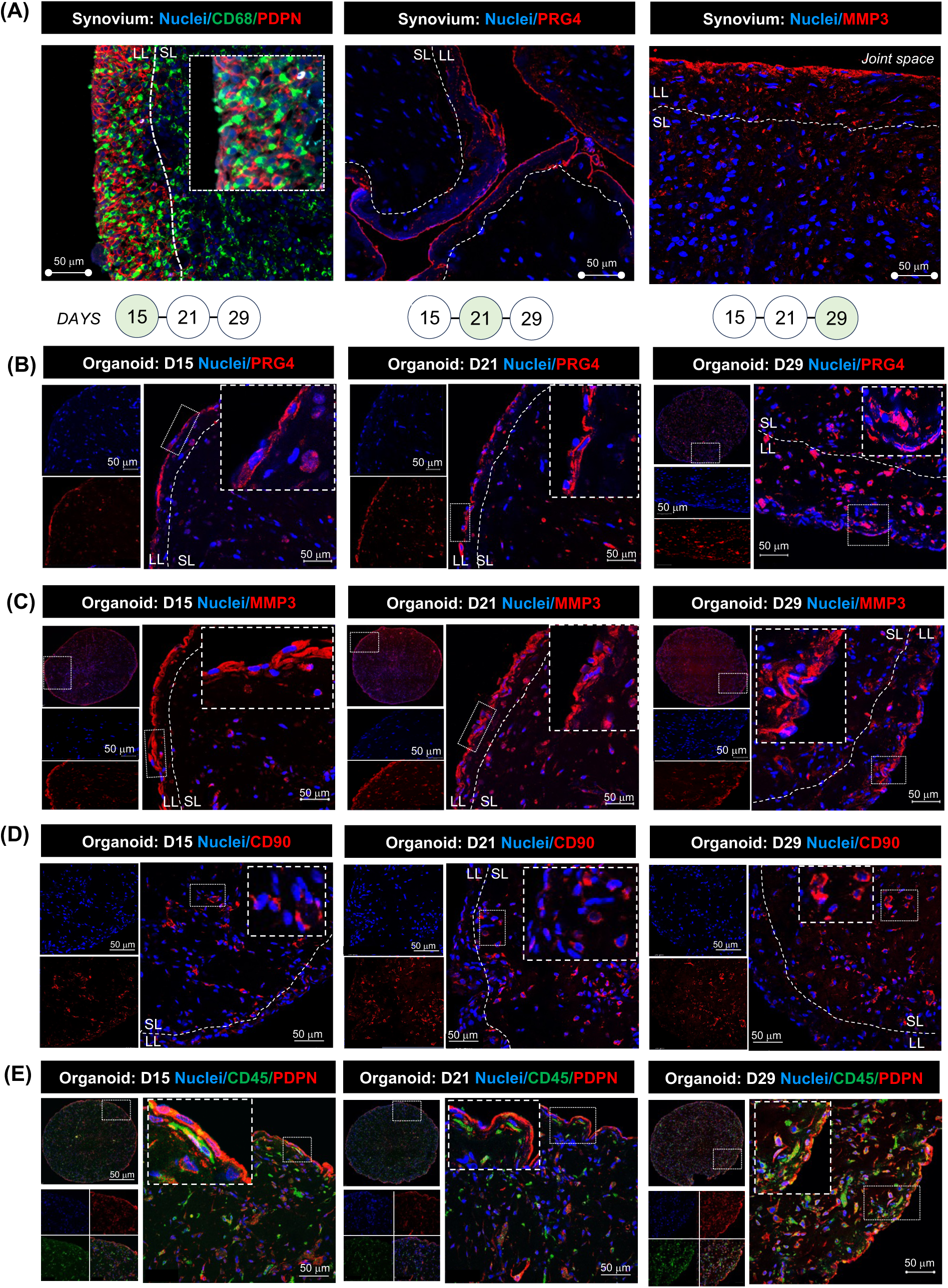
Myeloid-synovial stroma organoids recapitulate synovial tissue topography. **(A)** Representative confocal microscopy images (40×) showing immunofluorescence staining of the macrophage marker CD68 (green) and lining layer (LL) fibroblast markers PDPN, MMP3 (red), and PRG4 (red), with nuclei stained with DAPI (blue) in synovial tissue biopsy (ST). The inserts show enlarged cells in the selected regions of the tissue. Images are representative of ST biopsies from patients with active RA (n = 3–5). **(B–E)** Immunofluorescence staining of synovial organoids (SO) showing FLS with PRG4 (red) in (B), MMP3 (red) in (C), CD90 (red) in (D), and PDPN (red) in (E) as well as macrophages with CD45 staining (green) in (E) at different time points of SO development. Nuclei are stained with DAPI (blue) in all images. Data are representative of confocal microscopy images (40×). The inserts show enlarged cells in the selected regions of the SO. Data is representative of three independent experiments. Scale bars 50 µm for 40× magnification images. LL=lining layer, SL=sublining layer.

In summary, we have established myeloid–synovial stromal organoids that recapitulate the key features and anatomical organization of human synovial tissue.

### Synovial organoids support development of all identified STM subsets from monocytic precursors

Next, using the organoids, we investigated whether the synovium-like, spatially distributed stroma provides a permissive niche for the maturation of distinct synovial tissue macrophage subsets and their phenotypic clusters. First, we assessed whether the diversity of RA FLS is preserved in organoids. After digestion of 15-day old synovial organoids derived from stroma of ST biopsies from n=7 RA patients with active disease, FLS were identified by PDPN expression (lining FLS) and by combined PDPN and CD90 (THY1) expression (sublining FLS) **(Fig.4A).** Lining FLS accounted for 9.8± 2% (Mean SEM) and sublining FLS for 88± 1.5% of the total FLS population **(Fig.4B)**, closely resembling the ratio observed in synovial tissue taken ex vivo from patients with active RA (*6, 7*). All organoid FLS were subjected to single-cell transcriptomic analysis to establish their detailed phenotypic composition. The data were integrated with a reference single-cell dataset of RA synovium (n = 7) (*12*) using scANVI (*20*). UMAP visualization of the integrated dataset **(Fig. S4A)**, which was then separated into organoid and reference FLS **(Fig. 4C)**, together with prediction scores assessing the fit of organoid FLS to tissue FLS **(Fig. S4A)** demonstrated that organoids maintained the composition and phenotypes of active RA synovial tissue FLS. This confirmed the presence of lining PRG^pos^ FLS and further established that organoid sublining CD90 (THY1)^pos^ FLS encompass all previously described clusters (*7, 14*). These include perivascular CD34^pos^ and CXCL14^pos^GAS6^pos^ clusters, the proinflammatory HLA-DR^pos^CXCL12^pos^ cluster, which is the most expanded in active RA synovium (*6*), and the POSTN^pos^ collagen-producing cluster **(Fig. 4C and fig.S4B).** Furthermore, a time-course experiment **(fig. S4C)** confirmed that FLS phenotypes were preserved beyond day 15 in synovial organoids, highlighting their sustained potential to provide niche signals for immune cells. Thus, to determine whether these spatially and three-dimensionally organized FLS clusters provide cues for myeloid precursors to differentiate into distinct STMs, we performed single-cell transcriptomics of blood-derived myeloid precursors at baseline (day 0, prior to organoid embedding) and of their progeny in 15-day-old organoids (n = 7) **(Fig.4A).** Both datasets were then integrated with reference synovial tissue myeloid cells from active RA patients (n = 7). UMAP visualization of the integrated datasets **(Fig.4D and fig.S5A)**, together with prediction scores assessing the fit of organoid myeloid cells to synovial tissue myeloid cells **(fig.S5A)**, demonstrated that synovial organoids supported maturation of all synovial tissue macrophage populations and their phenotypic clusters identified in human synovium **(fig.S5B).**

**Figure 4.**
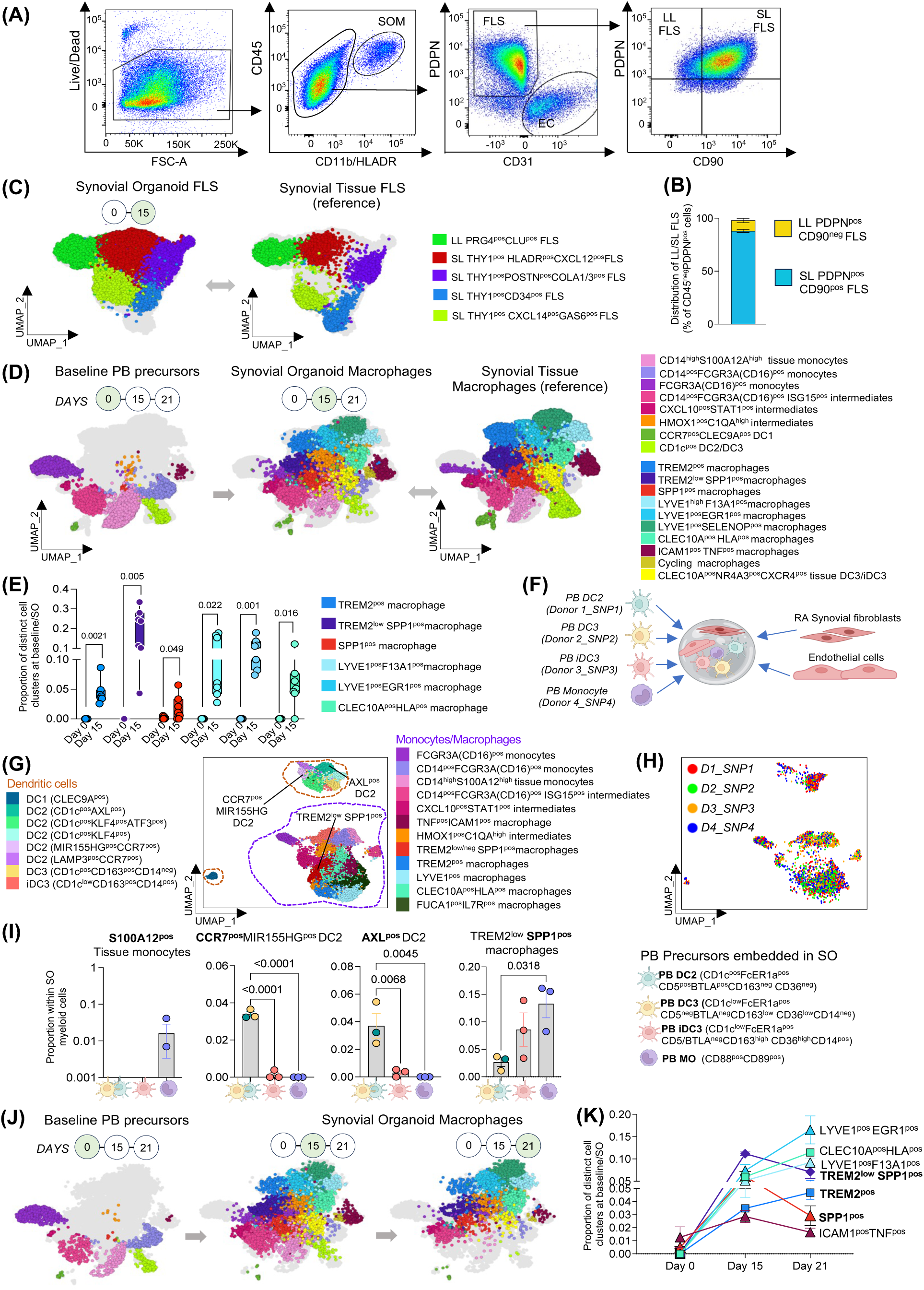
Human synovial organoids support development of synovial tissue macrophage clusters. **(A)** A representative gating strategy for the SO stromal and macrophage populations after digestion. **(B)** Flow cytometry validation of the frequency of lining layer (LL) and sublining layer (SL) FLS populations in synovial organoids (FLS from biopsies of n=3 RA patients with active disease, with 3–5 technical replicates). **(C)** UMAP visualization of integrated (scANVI) single-cell transcriptomics data of synovial organoid FLS and reference synovial tissue FLS. The dataset includes synovial fibroblasts from 15-day-old synovial organoids (31,101 cells) derived from synovial tissue biopsies of n=7 RA patients with active RA, and FLS directly sorted from synovial tissue (7,395 cells, n=7 Active RA tissues, reference). Data were collected across 14 independent experiments. Each dot represents an individual cell (42,179 cells in total). **(D)** UMAP visualization of integrated myeloid cell clusters from day 15 synovial organoids (11,349 cells) generated using FLS from biopsies of n = 7 patients with active RA, monocyte/DC populations from baseline peripheral blood samples (n = 7 healthy donors, including n=6 matching the 15-day organoids; 8,421 cells), and reference myeloid cells directly sorted from synovial tissue of 34 different RA patients with active disease (22,655 cells), across 14 independent experiments. **(E)** Proportional distribution of TREM2^pos^, TREM2_ˡ_ᵒʷSPP1^pos^, TREM2^neg^SPP1^pos^, FOLR2^high^LYVE1^pos^, FOLR2^high^EGR1^pos^, and FOLR2^high^CLEC10A^pos^ macrophage clusters at baseline (Day 0) and in synovial organoids (SO) on Day 15, based on n = 7 experiments as shown in (D). Data are presented as the median with the interquartile range. Each dot represents a patient derived SO (n=7). A paired T-test between matched baseline monocytes (Day 0) and matched myeloid cells at Day 15 organoids, with exact p-values displayed on the graph. SO=synovial organoids. **(F)** Schematic of SNP-based human synovial organoid system to deconvolute ST myeloid cell cluster differentiation trajectories. PB DC2, DC3, iDC3, and monocytes, each sorted from the blood of different healthy donors (with different SNP pattern), were embedded together in Matrigel with synovial biopsy-derived fibroblasts from active RA and endothelial cells (HUVEC). **(G)** UMAP visualizing scRNAseq data of CD45-positive cells (n=2273 cells) from SO as in (F) that was integrated with baseline PB DC2, DC3, iDC3 and monocytes (n=1646 cells) and with down-sampled reference scRNAseq data of RA synovial tissue myeloid cells (n=1915 cells). **(H)** UMAP as in (G) coloured by the cells from different donors based on a donor unique SNP pattern (Souporcell_v 2.4, as described in the methods). **(I)** Bar plots with SEM showing the frequency of different ST-DCs and macrophage clusters that developed from different blood precursors. One-way ANOVA with Dunnett’s corrections for multiple comparisons was used. The exact p-values are provided on the graphs. **(J)** UMAP visualization of integrated myeloid cells from organoid at day 15 and day 21 with matching baseline PB monocytes/DCs (n=2 of baseline and n=4 SO technical replicates). **(K)** Proportional distribution of TREM2^pos^, TREM2_ˡ_ᵒʷSPP1^pos^, TREM2^neg^SPP1^pos^, FOLR2^high^LYVE1^pos^, FOLR2^high^EGR1^pos^, and FOLR2^high^CLEC10A^pos^ macrophage clusters from (J) over time, comparing dynamic of their development from baseline monocytes to day 15 and day 21 synovial organoid macrophages. Data are presented as a connected Mean ± SEM of 4 technical replicates.

Moreover, integration of organoid- and tissue-derived myeloid cells with their blood-derived precursors (baseline) enabled us to trace their progression along the monocyte-to-macrophage differentiation trajectory and to assess the frequency of each cluster across healthy, active RA, and RA in remission joints **(Fig.1C and Fig.4D).** Organoid and tissue populations that clustered with blood cells were annotated as tissue monocytes, such as the alarmin producing S100A12^pos^ cluster characteristic of early naïve-to-treatment active RA **(Fig.1D and fig.S1-2)** (*7*). Populations that clustered with blood cells but, unlike their blood counterparts, expressed FOLR2 **(fig.S5C-D)**, a marker of tissue macrophages (*7*), were annotated as an early intermediate stage on the macrophage trajectory. This stage includes CXCL10^pos^STAT1^pos^ and ISG15^pos^ populations, which are rare in healthy individuals but increase in abundance and cell proportion in patients with active disease, both treatment-naïve and treatment-resistant **(Fig.1D and fig.S1-2)** and resolved in sustained remission (*7, 12, 14, 17, 21*).

Populations that clustered with both blood and synovial tissue macrophages, were annotated as a later intermediate stage and included HMOX1^pos^C1QA^pos^ FOLR2^pos^ cluster that was also increased in abundance particularly in naïve-to-treatment RA as compared to healthy and remission RA **(Fig.1D and fig.S1).** The populations that were absent in blood (baseline) and emerged only in synovial organoids and synovial tissue, were annotated as macrophages. Quantitative evaluation of synovial tissue macrophage clusters in organoids (n = 7) at day 15 **(Fig.4E)** revealed the consistent emergence of synovial tissue–specific resident lining layer TREM2^pos^ macrophages, with the highest frequency observed in their pathogenic TREM2^low^SPP1^pos^ phenotype, reflecting the lining layer composition characteristic of active RA (*7*) (**Fig.1D and fig.S1-2).** Synovial tissue organoids also supported the development of distinct phenotypic clusters of tissue-resident perivascular LYVE1^pos^ STMs. These included ERG1^pos^ and SELENOP^pos^ clusters **(Fig.4E)**, each exhibiting distinct pro-inflammatory phenotypes: CCL3/CCL4 expression in ERG1^pos^ STMs, and CXCL9/CCL18 expression in SELENOP^pos^ STMs **(fig.S5B)**. These phenotypes recapitulate those abundant in early, treatment-naïve active RA synovium compared with healthy tissue and RA in remission **(Fig.1D, fig.S1-2).** In addition, synovial tissue organoids supported the development of tissue-resident antigen-presenting CLEC10A^pos^HLA^pos^ macrophages, uniquely characterized by the highest expression of antigen-presenting pathways among all STMs, as well as the emergence of infiltrating pathogenic FCN1^pos^SPP1^pos^ and ICAM1^pos^TNF^pos^ subsets **(Fig.4E)**, all increased in abundance and proportion in active RA synovium as compared to healthy and remission RA **(Fig.1D).**

Single-cell analysis of blood monocyte precursors at baseline revealed low-level contamination with conventional (c) DC2, DC3 (12 ± 8%, mean ± SEM) and iDC3 (0.3 ± 0.2%). Accordingly, by day 15, organoids had developed resident synovial tissue–associated cCD1c^pos^ DC2/DC3 phenotypes (0.4 ± 0.2%) and, notably, an RA-associated iDC3 phenotype (SDS^pos^NR4A3^pos^CXCR4^pos^) (2 ± 1%) **(Fig. 4D)**, consistent with DC profiles observed in active RA synovium **(Fig.1D)** (*12*). To confirm that only monocytes give rise to lining layer STMs and to directly compare the differentiation potential of different peripheral blood myeloid precursors into distinct tissue myeloid cell clusters within organoid niches, we sorted monocytes, as well as PB conventional DC2, DC3, and iDC3, from four healthy donors using a recently established multi-marker sorting strategy specific for each precursor **(fig.S6)** (*12, 19*), and embedded them in synovial organoids. Each organoid contained all four PB subsets, with each subset derived from a different donor to allow deconvolution of their differentiation outcomes based on donor-specific SNP patterns **(Fig.4F).** The immune compartment of the organoids was then analysed by single-cell transcriptomics, and the data integrated with a reference myeloid synovial tissue scRNAseq dataset **(Fig.4G)** as in Fig.4D. The developmental trajectories of annotated clusters were deconvoluted based on donor-specific SNPs **(Fig.4H).** This analysis revealed that only monocytes, and not DC2/3, gave rise to the tissue S100A12^pos^ population, whereas only DC2 and DC3 gave rise to specific synovial tissue cDC2 clusters, namely AXL^pos^ DC2 and CCR7^pos^ mReg DC, as recently described, validating our system (*12*), **(Fig.4I).** Deconvolution of the trajectory of active RA lining layer TREM2^pos^SPP1^pos^ cells in this multi-precursor, SNP-based cell fate–tracking organoid system confirmed that they differentiate from blood monocytes not cDC2/3. It also revealed that blood iDC3, which share a transcriptomic profile with monocytes (*12*), likewise have the potential to differentiate into TREM2^pos^SPP1^pos^ macrophages. Tracking the transition from baseline monocytes to day 15 and then day 21 organoid myeloid cells revealed a time-dependent differentiation trajectory, in which tissue monocytes such as the S100A12^pos^ population progressively disappeared, while the monocyte-to-macrophage intermediate state (HMOX1^pos^C1QA^high^ cluster) and synovial tissue–resident macrophage populations, including lining TREM2^pos^, perivascular LYVE1^pos^, and antigen-presenting CLEC10A^pos^ clusters, increased over time **(Fig. 4J–K and fig. S7).**

These data show that spatially organized RA synovial stroma supports the survival of blood-derived myeloid precursors and directs their differentiation into the full spectrum of STM, including both homeostatic and pathogenic subsets and clusters.

### Synovial stroma drives development of lining layer TREM2^pos^ STMs and its pathogenic TREM2^low^SPP1^pos^ phenotype that is augmented by synovial fluid

Finally, we sought to validate the scRNAseq findings at both functional and spatial levels. We focused specifically on TREM2^pos^ STM clusters, given their critical role in joint homeostasis and pathology (*5, 7*). Multiparameter flow cytometry of digested synovial organoids at day 15, with matched baseline blood precursors as comparators, confirmed high expression of the macrophage markers FOLR2 and HLA-DR in CD45⁺ organoid cells **(fig.S8A).** Approximately 40% of these cells were identified as TREM2^pos^ lining layer macrophages, of which ∼10% produced SPP1 protein, consistent with a TREM2^pos^SPP1^pos^ lining layer STM phenotype. As expected, none of these phenotypes were detected at baseline **(Fig.5A–B).** Immunofluorescence staining of synovial organoids further confirmed that TREM2 and SPP1 expressing macrophages predominantly localized to the lining layer **(Fig.5C–D),** whereas LYVE1^pos^ macrophages marked the perivascular sublining niche adjacent to tube-like endothelial structures **(Fig.5E–F).** Altogether, these findings provide further evidence that synovial stroma drives the differentiation and spatial localisation of lining layer TREM2^pos^ macrophages and promotes their pathogenic SPP1-producing cluster.

**Figure 5.**
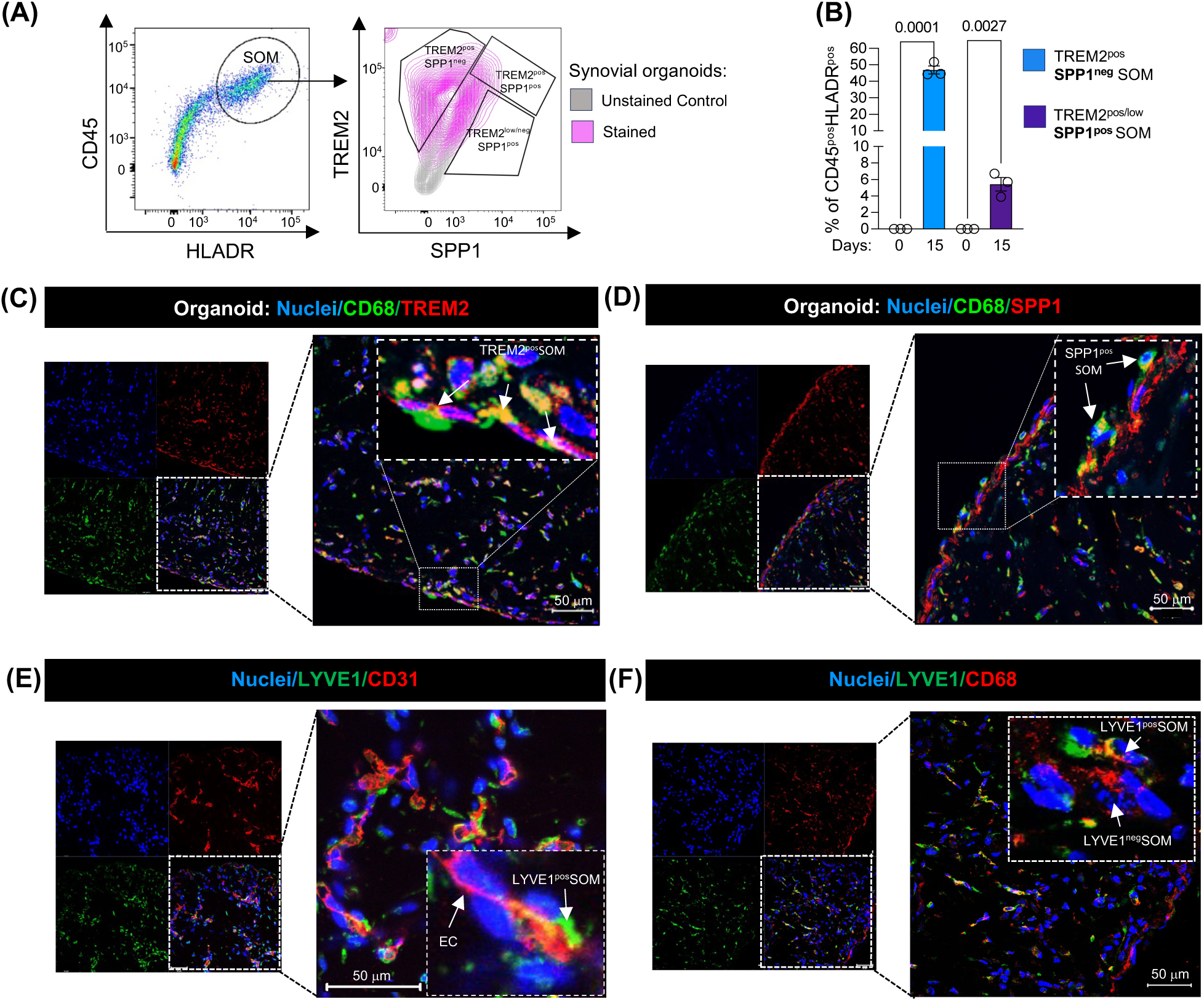
Human synovial organoids support development of LL TREM2^pos^, its pathogenic SPP1^pos^ phenotype and SL LYVE1^pos^ STMs. **(A)** Representative FACS plots showing synovial organoid (SO) macrophage (SOM) gated based on CD45 and HLADR expression at day 15 SO. (B) The frequency of TREM2^pos^SPP1^neg^ and TREM2^pos/low^SPP1^pos^ macrophages within CD45^pos^HLADR^pos^ myeloid cells at the baseline (day 0) and day 15 of SO of active RA. Data are represented as Mean ± SEM from 3 biological replicates (FLS from n=3 patients with active RA). **(C-D)** Immunofluorescence images showing localisation of TREM2^pos^ (C) and SPP1^pos^ macrophages (D), CD68, green; TREM2 and SPP1, red; and Nuclei, blue. **(E-F)** Immunofluorescence images showing presence of LYVE1^pos^ (green) macrophages near CD31-positive (red) endothelial cell (EC) tube-like structures. (C-F) Data shows representative confocal microscopy images (40x) of SO generated from biopsies of n=3-5 RA patients with active disease in three independent experiments. Scale bars, 50µm. SOM=synovial organoid macrophages. Lining layer, LL; Sublining layer, SL.

Because lining layer TREM2^pos^ STMs are directly exposed to synovial fluid (SF), which in active RA contains, in addition to lubricants, pro-inflammatory mediators released by inflamed tissue, we next assessed its impact on the lining layer STM phenotype. Synovial organoids were exposed to cell-free 10% synovial fluid, pooled from five RA patients, during the last seven days of culture, followed by the analysis of synovial organoid myeloid cell phenotypes by single-cell transcriptomics **(Fig.6A–B).** This intervention selectively increased the frequency of SPP1-expressing lining layer STMs **(Fig. 6C–D).** Multiparametric flow cytometry **(Fig.6E)** revealed an SF-induced shift in STMs from homeostatic TREM2^pos^SPP1^neg^ toward RA associated TREM2^neg^ SPP1 protein producing macrophages **(Fig.6F–G).** Immunofluorescence staining of synovial organoids exposed to synovial fluid confirmed the loss of the TREM2^high^ phenotype and the expansion of SPP1-producing macrophages within the hyperplastic lining layer niche **(Fig.6H–I and fig.S8B-C**), closely recapitulating active RA tissue **(Fig.1F).** Together, these findings demonstrate that the disorganized and hyperplastic lining layer in active RA, enriched in pathogenic TREM2^pos^SPP1^pos^ and TREM2^neg^SPP1^pos^ macrophages, is orchestrated by signals from both from both adjacent synovial fibroblasts and synovial fluid.

**Figure 6.**
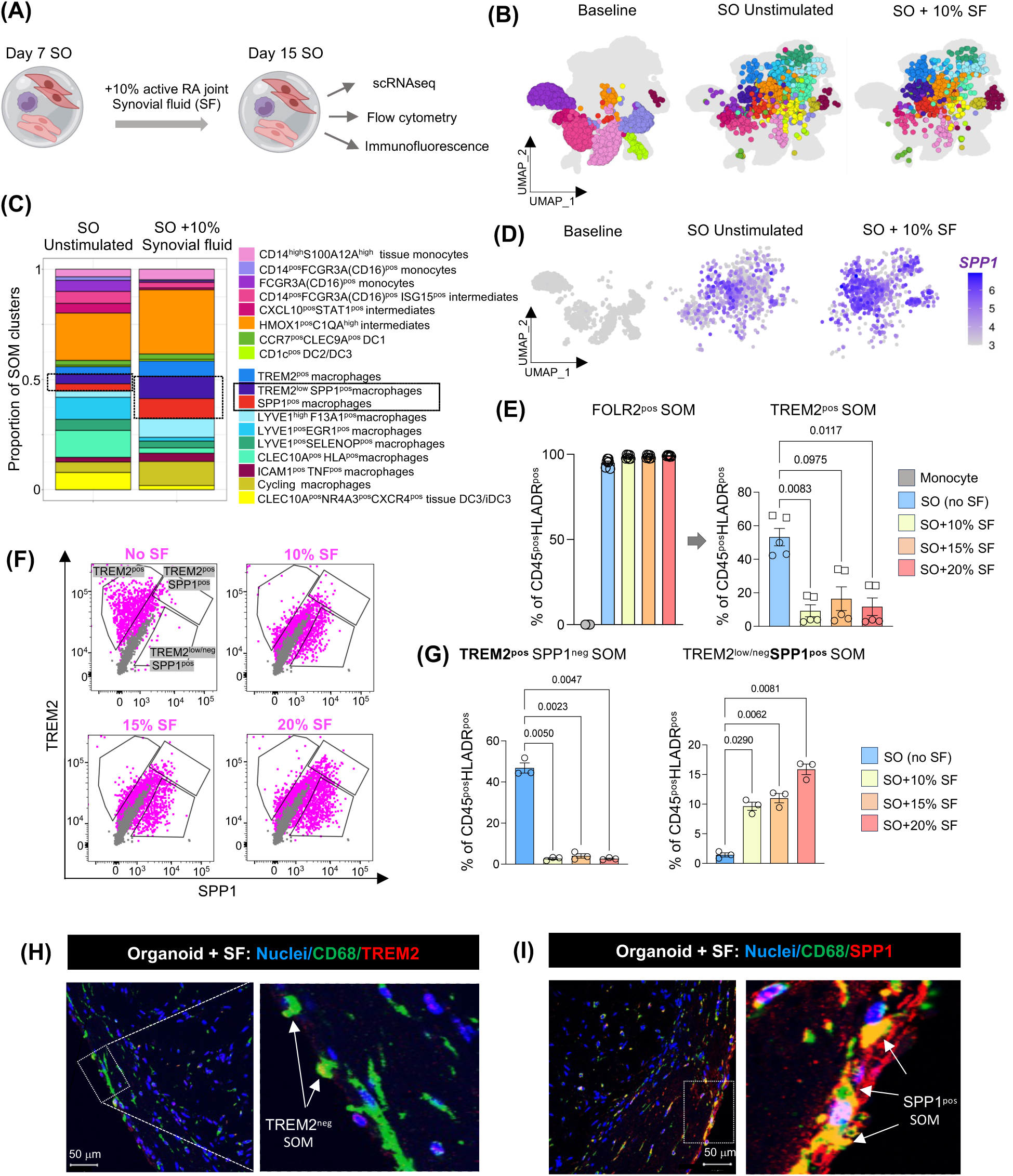
Synovial stroma drive development of lining layer TREM2^pos^ STMs and its pathogenic TREM2^low^SPP1^pos^ phenotype that is augmented by inflamed synovial fluid. **(A)** SO were generated as in Figure 2C. To mimic acute inflammatory stimulation, at day 7, 10% of cell-free synovial fluids (SF) isolated from active RA joints were added to SO for subsequent 7 days. **(B)** UMAP visualization of single cell transcriptomics of myeloid cells from synovial organoids cultured with (985 cells) or without 10% synovial fluid (1,160 cells). **(C)** Proportional distribution of myeloid cell clusters identified by scRNAseq, comparing organoids cultured with and without SF. **(D)** UMAP feature plot showing the expression levels of SPP1 in myeloid cells at baseline and from synovial organoids (SO) cultured with or without 10% SF. **(E)** Flow cytometry data showing the percentage of FOLR2 and TREM2 positive macrophages within CD45^pos^HLADR^pos^ myeloid cells in synovial organoids cultured with 0%, 10%, 15%, or 20% SF. Expression of FOLR2 in baseline monocytes is also shown while TREM2 on monocytes is shown in Figure 5B. Data is presented as a bar plot showing Mean ± SEM of 2 biological replicates (FLS derived from biopsies of 2 RA patients with active disease) with two or three technical replicates each. Each FLS donor represented by different symbol. Kruskal-Wallis test with Dunn’s corrections for multiple comparison with exact p values on the graphs. **(F)** Representative dot plot showing surface and intracellular expression of TREM2 and SPP1 in macrophages derived from SO cultured in the absence or presence of different concentration of SF. Macrophages were gated as HLADR and FOLR2 positive. **(G)** Flow cytometry evaluation of frequency of TREM2^pos^SPP1^neg^ and TREM2^low/neg^SPP1^pos^ SOM in SO at day 15. Data is presented as a bar plot with Mean ± SEM of three technical replicates. **(H-I)** Representative immunofluorescence staining of TREM2 (H) and SPP1 (I) positive macrophages (CD68) in SO in the presence of 20% SF. CD68 (green) and Nuclei DAPI (blue) in H-I, TREM2 (red) in H and SPP1 (red) in I. Representative confocal microscopy images (40× magnification) of SO generated with FLS from biopsies of three active RA patients, obtained from three independent experiments. Scale bars, 50µm. Synovial fluid, SF pooled from n=5 RA patients with active diseases. SOM=synovial organoid macrophages.

## DISCUSSION

In this study, we defined the origins of human synovial tissue macrophage subsets, identified the tissue niches that shape their functional phenotypes, and established a model that faithfully recapitulates human synovial niches to enable exploration of their therapeutic potential. Using SNP-based cell fate–tracking approaches, both in a patient who received heterologous bone marrow transplants and in ex vivo in synovial organoid models, we demonstrated that all human STMs, including tissue-resident subsets, can originate from blood monocytes. Insights from human embryonic joints show that STMs appear only after the synovial stroma acquires a joint-specific phenotype (e.g., PRG4^pos^ lining FLS), suggesting that the synovial stroma establishes an initial tissue niche in the joint that then provides myeloid precursors with joint-specific cues. Human synovial organoids confirmed that three-dimensionally organized synovial tissue stroma drives the differentiation of myeloid precursors into distinct, anatomically restricted STM subsets located within spatially and transcriptionally defined niches. These include key subsets such as TREM2^pos^ STMs in the lining and LYVE1^pos^ STMs in the sublining.

In addition, an extended single-cell synovial tissue myeloid atlas across health and RA disease trajectories, combined with tissue spatial mapping and validation in synovial organoids, provided novel insights into the macrophage phenotypes in the hyperplastic lining layer of active RA. We uncovered that the TREM2^low^ cluster that replaces homeostatic TREM2^pos^ cells in the RA lining layer specifically produces high levels of pathogenic SPP1, which drives neutrophil and infiltrating monocyte activation (*17*). Our recent findings also indicate that SPP1^pos^ macrophages increase sensory neuron firing, potentially activating nociceptors at the lining–sublining interface and thereby contributing to pain (*2, 22*). Together with emerging data from children with early-diagnosed juvenile idiopathic arthritis (*11*), patients with pre-clinical RA (*15*), and mouse models (*23*), these findings suggest that the pathogenic transition from TREM2^pos^ to TREM2^low^SPP1^pos^ macrophages in the lining may be responsible for creating an inflammation, autoimmunity-, and pain-permissive niche that enables establishment of pathogenic immune response in the joints (*2, 5, 22, 23*). Synovial organoids demonstrated that this pathogenic phenotype of lining STMs is driven by RA stromal niche signals and further amplified by inflamed synovial fluid.

We previously uncovered the key role of endothelial cells in establishing the sublining stromal niche in the RA synovium (*24*). Here, we demonstrate that this niche provides in turn unique signals for the development of perivascular LYVE1^pos^ STMs, particularly clusters with pro-inflammatory phenotypes, such as CCL3/CCL4-expressing ERG1^pos^ and CXCL9 and CCL18-expressing SELENOP^pos^, which are features of active RA synovium compared with healthy tissue. Through reciprocal interactions with their perivascular niche, these cells promote immune cell infiltration into the tissue, likely facilitating the localization of the immune response within the joint (*7, 13*).

Adult fibroblasts maintain a lasting imprinting of their organ of residence and position, reflecting their embryonic origins. This is orchestrated by HOX genes, tissue-specific transcription factors, and epigenetic imprinting, which together help preserve the identity of each organ’s fibroblasts (*25, 26*). This organ-specific stroma support myeloid cells by providing positional cues through the extracellular matrix, pan-macrophage factors (e.g., CSF1, IL-34, CSF2), and tissue-specific signals that guide their differentiation into organ- and niche-specialized subsets (*3, 25–27*). Emerging evidence suggests that aging, non-physiological loading, trauma, and chronic inflammation, such as in rheumatoid arthritis, can induce joint-specific epigenetic changes in the stroma that persist as inflammatory tissue memory (*28–30*). Our findings indicate that these inflammation-imprinted stromal cells provide altered niche signals to myeloid cells, fostering the local maturation of pathogenic STM phenotypes in RA. The tractable ex vivo myeloid–synovial stromal organoid system established in this study provides a platform to define those stromal-derived cues that drive the differentiation of distinct STM clusters and to explore therapeutic strategies for resetting the synovial stroma to reinstate homeostatic tissue signals—for example, to shift the lining-layer STM phenotype from an SPP1-producing to a regulatory VSIG4^pos^TREM2^pos^ state.

*Limitations of the study.* We cannot exclude the possibility that treatments used for bone marrow ablation also eliminated any in situ proliferating fetal STM precursors, leaving an empty tissue niche primed for repopulation. Nevertheless, our study demonstrates that blood-derived monocytes are capable of differentiating into all human STM clusters and, together with synovial fibroblasts, establish distinct pro-inflammatory or pro-resolving synovial niches.

## MATERIALS AND METHODS

### Study Design

The objective of this study was to define the origin and developmental trajectory of human synovial tissue macrophages (STMs), with a particular focus on the phenotypes of the lining layer, and their abundance between different joint conditions. This was investigated by: **(i)** Examining the origin of STMs in the synovium of a patient that underwent heterologous bone marrow transplantation; **(ii)** Establishing a myeloid–stromal organoid system to enable tracking of STM development; **(iii)** Assessing the abundance and phenotypes of different STM populations in different joint conditions using our extended peripheral blood and synovial tissue (PB/ST) myeloid cell atlas (*12*), supplemented with 16 additional synovial tissue biopsies collected for this study, mounting to total n=11 healthy, n=23 active RA, and n=11 in sustained disease remission), and **(iv)** Investigating the development of the synovial niche in human embryonic joints. **Approach: (i)** We recruited a 59-year-old male patient with arthralgia, who had undergone a heterologous bone marrow transplant four years earlier for acute nonlymphocytic leukemia. He underwent synovial tissue biopsies and matched peripheral blood collection. Comprehensive Ultrasound assessment was conducted that showed a lack of subclinical inflammation (absence of Power Doppler signal). Previous treatments used for prevention of graft versus host disease were mycophenolate mofetil, cyclophosphamide and cyclosporin A, all stopped at least 2 years before the synovial tissue biopsy performance. **(ii)** To develop the synovial organoid system, 10 patients fulfilling the EULAR classification criteria revised criteria for RA (*31*) were enrolled and underwent ultrasound-guided ST biopsy of the knee as a part of ongoing recruitment to the SYNGem cohort (*7, 12, 32, 33*) (standard-of-care protocol, Division of Rheumatology and Clinical Immunology, Fondazione Policlinico Universitario A. Gemelli IRCCS – Università Cattolica del Sacro Cuore, Rome). RA patients were stratified into treatment-naïve and treatment-resistant RA (inadequate responders to conventional or biological Disease Modifying Anti-Rheumatic Drugs, c/b DMARDs) (*34*). The clinical and laboratory evaluation of each RA patient enrolled included DAS based on the number of tender and swollen joints of 44 or 28 examined, plus the erythrocyte sedimentation rate (ESR) and plasma C-reactive protein (CRP). Peripheral blood samples were tested for IgA-RF and IgM-RF (Orgentec Diagnostika, Bouty-UK), and ACPA (Menarini Diagnostics-Italy) using commercial Enzyme-Linked Immunosorbent Assay (ELISA) and ChemiLuminescence Immunoassay (CLIA) respectively. Demographic, clinical, and immunological features of this cohort are shown in **Table S1**. Healthy controls (n=12) that provided myeloid precursors for synovial organoids were recruited at University of Glasgow. **(iii)** Synovial biopsies were obtained from additional controls and RA patients, including individuals undergoing arthroscopy for meniscal tears or cruciate ligament damage with macroscopically and MRI-confirmed normal synovium (controls, n = 7), as well as from RA patients with active disease (n = 4) and those in sustained remission (n = 5), as described in (i). The demographic, clinical, and immunological characteristics of this cohort are presented in **Table S2**, and MacDonald et al (*12*). (**iv)** To investigate maturation of lining layer, human embryonic joints were obtained from elective terminations at 9-, 10-, 13- and 14-weeks post-conception and preserved for whole-mount immunostaining as described previously (*18*).

### Ethics

The study protocols on adult RA patients’ and healthy donors’ synovium and blood were approved by the Ethics Committee of the Università Cattolica del Sacro Cuore (28485/18, 30973/19 and 14996/20) and by the West of Scotland Research Ethics Committee (19/WS/0111 for healthy donors). Lower limb knee joints from foetuses were taken by the Human Developmental Biology Resource (HDBR) team post-medical or -surgical pregnancy termination. Protocols for collection and use of developmental joint samples were approved by the London-Fulham Research Ethics Committee under code 18/LO/0822. All samples were taken with written and informed consent obtained from all sample donors. A synovial fluid stock was made by pooling synovial fluid of RA patients with actively inflamed joints (n=5). Patient consent was obtained by all participants in written format according to the Declaration of Helsinki and the study was approved by the medical ethics committee of the Academic Medical Centre, University of Amsterdam, Amsterdam, the Netherlands.

### Generation of bone marrow transplant synovial tissue and peripheral blood single-cell dataset, and its integration with RA and healthy synovial tissue and blood datasets

Single cell transcriptomics (whole transcriptome) of knee synovial tissue biopsy and matched peripheral blood from a patient who underwent bone marrow transplant were performed using BD-Rhapsody platform as we described previously (*12*). Briefly, live cells were labelled with unique sample identifier tags (using the BD Human Single-Cell Sample Multiplexing Kit (633781/BD Bioscience) according to the manufacturer’s protocol. Cells were then loaded onto the scRNAseq BD Rhapsody Cartridge using the BD Rhapsody Cartridge Reagent Kit (633731) according to the manufacturer’s protocol. Single-cell cDNA was prepared using the BD Rhapsody cDNA Kit (633773). This was followed by single-cell Tag library preparation kit (633774) and mRNA library preparation with BD Rhapsody WTA (633801). Libraries were sequenced using Illumina NextSeq 500 (Glasgow Polyomics). Raw data analysis QC & Filtering were performed using the Seurat package (4.0.3) in R. Integration with our human synovial tissue atlas was then performed using the Seurat wrapper function (RunHarmony, SeuratWrappers, 0.3.0) for Harmony (*35*). Deconvolution of the hashtag/sample information (bone marrow donor versus bone marrow recipient) was performed using Souporcell (2.5) based on distinct donor and recipient SNPs information. Detailed descriptions of the integration and deconvolution methods are provided in our previous publication (*12*).

### Preparation Poly-HEMA coated plates for synovial organoids

The coating protocol was adapted from Wei et al (*24*). Twelve-well cell culture plates were coated with Poly-HEMA (Sigma-Aldrich, 192066-10G). To prepare the coating solution, Poly-HEMA was dissolved in ethanol (0.1 g/mL) preheated in a 60°C water bath. The solution was immediately vortexed at maximum speed for 2 minutes, followed by incubation at 60°C water bath for 10 minutes. Since Poly-HEMA crystals do not dissolve easily, the vortexing (2 minutes) and heating (10 minutes) steps were repeated until the granules had completely dissolved, which typically required 2–3 hours. Under sterile condition, the Poly-HEMA solution was aliquoted into 12-well plates at 1 mL per well, allowing for the plating of one SO per well. The plates containing the Poly-HEMA solution were placed in an oven at 60°C for 16–24 hours, allowing the ethanol to evaporate and leaving a thin plastic coating at the bottom of each well. Coated plates were stored at room temperature for up to one month.

### Generation of synovial organoids

Creating the synovial tissue organoid structure was conducted using three types of cells: early passage of synovial fibroblasts (passage number 1-3) of obtained from active RA synovial tissue biopsies (Table S1). After enzymatic digestion (*7*), synovial fibroblasts were cultured in DMEM (cat No 41965-039) supplemented with 10% foetal bovine serum, 100U/mL penicillin/streptomycin, 2mM Glutamax, 1% essential amino acid and 1% sodium pyruvate. Human umbilical vein endothelial cells (HUVECs) (passage number 2-8, cat No. C2519A, Lonza) were expanded in the presence of growth factors media (EGM-2, Lonza, cat No. CC-5035). Healthy blood monocytes were sorted based on their high CD88/89 expression. Briefly, live cells were gated, and lineage-positive cells (CD3, CD19, CD15, CD117, CD56) were excluded, while HLA-DR-positive cells were retained **(Table S3).** Monocytes were gated based on CD88 and CD89 expression. Individual organoid was created using 200 x10^3^ synovial fibroblasts combined with 200 x10^3^ HUVECs and 100 x10^3^ (CD88^pos^CD89^pos^ monocytes) in a single droplet of Matrigel (Matrigel® Growth Factor Reduced Basement Membrane Matrix, Phenol Red-Free, Corning, cat No 356231). Organoids were seeded on polyHEMA coated plates that were pre-warmed at least for one hour in a 37°C incubator. After one hour incubation, organoids were supplemented with 2ml EGM-2 media minus hydrocortisone and incubated for various time points (day 7, 15, day 21 and day 29), in the 37°C, 5% CO_2_ incubator. Every 3–4 days, 1.5 ml of medium was carefully collected from each well and replaced with 2 ml of fresh medium. In some experiments, on day 7 after SO generation, the organoids were cultured in the absence or presence of 10%, 15%, or 20% synovial fluid for varying durations (until days 15, 21, or 29).

### SNP-based human synovial organoid system

Peripheral blood (PB) DC2, DC3, iDC3, and monocytes were sorted from the PB of four healthy donors using the gating strategy recently described (*12*) and presented in **Figure S6.** Briefly, live cells were gated, and lineage-positive cells (CD3, CD19, CD15, CD117, CD56) were excluded, while HLA-DR-positive cells were retained. Monocytes were gated based on CD88 and CD89 expression. Plasmacytoid DCs (pDCs) and pre-DCs were excluded based on CD123 expression, while DC1 cells were excluded based on CD141 expression. The remaining myeloid DCs were gated based on CD1c and FcεRIα expression. DC2 cells were sorted based on CD5 and BTLA expression, while DC3 and inflammatory DC3 (iDC3) were distinguished by their differential expression of CD14 and CD36. Detailed on antibodies used are provided in **Table S4.** Four sorted populations (monocytes, DC2, DC3 and iDC3)—each from a different donor—were mixed in equal numbers (50,000 cells each) to generate a complete organoid, as described above. After 15 days, the organoids were digested, CD45⁺ cells were sorted, hash-tagged using BD Human Single-Cell Sample Multiplexing Kit (633781/BD Bioscience, and subjected to scRNAseq using the BD Rhapsody system, as detailed above. Souporcell (v2.4) was used to determine the SNP genotype of each donor at baseline and of each myeloid cell within the organoids after two weeks (*36*). Picard GenotypeConcordance (v2.18.29) was then used to match each cell’s SNP profile to its corresponding healthy donor, thereby tracing the origin of the myeloid cells back to their blood precursors.

### Live imaging of synovial organoids

PB monocytes were pre-labelled with cell tracker (C34567, Invitrogen, Excitation 549/15 nm, Emission: 630/90nm) while fibroblasts were labelled with cell tracker CFSE (C34554, Invitrogen, Excitation 452/45 nm, Emission: 512/23 nm) and combined with unlabelled HUVECs in a single droplet of Matrigel at day 0. After 48h, the continuous live recording was conducted using Omni imaging system for live imaging over 29 days. EGM-2 media (minus hydrocortisone) was replenished every 3-4 days.

### Optimization of Myeloid Cell Number for Synovial Organoid Generation

To determine the optimal number of immune cells for SO generation, we performed titration experiments using varying cell densities ranging from 0 to 150 × 10³ cells per organoid. SOs were digested at 7, 14, and 21 days to assess cell number, composition, and viability. Each time-point was evaluated with two or more replicates per time-point, resulting in a total of 20 SOs analyzed.

### Immunohistochemistry and Immunofluorescent staining of synovial organoids and reference RA synovial tissue biopsies

Synovial organoids were rinsed twice by immersion in 5 ml of PBS at room temperature in a 6-well plate. They were then fixed in 1.5 ml of 4% paraformaldehyde (PFA) at room temperature and stored overnight at 4 °C. The following day, the organoids were washed twice with PBS and dehydrated with 2 ml of freshly prepared 70% ethanol, as previously described (*24*). SOs were then embedded in paraffin and serially sectioned at 8 μm thickness. All staining procedures were performed at room temperature using primary antibodies and species-specific fluorophore-conjugated secondary antibodies. Details of the antibodies are provided in **Table S5.** Tissue sections were heated in a tissue-drying oven (GenLab) for ∼30 min at 60 °C, then deparaffinized twice in 100% xylene for 5 min each. Sections were rehydrated twice through a descending ethanol series (100%, 90%, and 70%) for 3 min each. For antigen retrieval, slides were immersed in either 0.01 M citrate buffer (pH 6.0; HDS05-100, TCS) or Dako Target Retrieval Buffer (pH 9.0; S2367), depending on the antibody, and heated in a microwave at 50% power for 4 min, followed by 30% power for 6 min. Slides were cooled for 15 min, washed in distilled water for 5 min, and rinsed twice in TBST (TBS with 0.025% Triton X-100; 93443, Sigma) for 5 min each. Sections were then blocked for 1 h at room temperature in 10% normal human serum (ready-to-use; 31876, ThermoFisher) with 1% BSA (A2153, Sigma) in TBS, supplemented with 10% normal serum from the host species of the secondary antibody (e.g., goat serum; 31872, ThermoFisher) to prevent non-specific binding, in a humidified chamber. To assess Collagen Type I (polyclonal) and Collagen Type III (polyclonal) in SO sections, slides were incubated overnight with primary antibodies or appropriate isotype controls. The following day, sections were washed and incubated with MP-7401/ImmPRESS HRP Horse Anti-Rabbit secondary antibody for 30 min. After two washes in TBST (5 min each), staining was developed using DAB chromogen according to the manufacturer’s instructions. Sections were then washed in water for 5 min, counterstained with Harris hematoxylin for 15 s, rinsed under running water for 2 min, and dehydrated through an ascending alcohol series. Finally, sections were mounted with DPX mounting medium, air-dried, and covered with coverslips. Images were acquired using an Olympus BX41 microscope with a DP25 camera and Axiovision software. *Immunofluorescent staining*. To assess the topography of SO, sections were incubated overnight at 4 °C with primary antibodies against the following markers: PDPN (polyclonal), CD90 (clone 7E1B11), PRG4 (clone 9G3), MMP3 (monoclonal, EP1186Y), CD31 (clone EPR3094), TREM2 (polyclonal), and SPP1 (polyclonal), either alone or in combination with antibodies against CD45 (monoclonal, IgG2b) or CD68 (clone PG-M1), or with appropriate isotype controls, all diluted in the blocking buffer described above (**Table S5**). The following day, sections were washed twice in TBST for 5 min each and incubated with appropriate secondary antibodies in dilution buffer (TBS with 1% BSA) for 1 h at room temperature. After incubation, sections were washed twice in TBS (5 min each) and counterstained with VECTASHIELD® Vibrance™ Antifade Mounting Medium with DAPI (H-1800-2). Sections were visualized using a Zeiss LSM 880 confocal microscope with a water immersion LD C-Apochromat ×40/1.3 NA objective, and images were acquired using Zen Black software (Zeiss). Image processing, including brightness/contrast adjustment and background subtraction, was performed using the same software. *RNAscope staining.* For detection of SPP1 transcripts in synovial tissue macrophages, we used the RNAscope Multiplex Fluorescent v2 assay (Advanced Cell Diagnostics, Newark, CA, USA, cat. no. 323100) on formalin-fixed paraffin-embedded (FFPE) synovial tissue sections. Tissue blocks were sectioned at 4–5 μm, deparaffinized, and processed for target retrieval and protease treatment according to the manufacturer’s instructions. The SPP1 probe (cat. no. 420101) and VSIG4 probe (cat. no. 446361) were hybridized to the sections. Following in situ hybridization, sections were immunostained with Alexa Fluor® 594 anti-CD68 antibody (clone KP1; cat. no. ab282650, Abcam) to identify macrophages. Nuclei were counterstained with SYTO™ 83 Orange Fluorescent Nucleic Acid Stain (ThermoFisher, S11364)

### Immunofluorescent Staining of Human Embryonic and Adult Joints and Multiplexed Image Acquisition

*Tissue Embedding and Epitope Retrieval*. Developmental joint sections were fixed in 10% neutral-buffered formalin for at least 48 hours prior to paraffin embedding. Tissue sections were cut at 6 μm for developmental joints and 5 μm for adult joints. Sections were baked overnight at 60 °C, then dewaxed in xylene and rehydrated through a descending ethanol gradient from 100% to 50%. Slides were permeabilized with 0.3% Triton X-100 (Invitrogen) and washed with PBS. Heat-induced epitope retrieval was performed in a pressure decloaking chamber at 110 °C for 20 min at 6.1 PSI using two buffers: pH 6.0 citrate (Sigma, C9999) and pH 9.0 Tris-based solution (Agilent, S2367). After cooling, slides were blocked overnight at 4 °C in 10% donkey serum and 3% BSA. *Immunofluorescent Slide Staining.* Following blocking, slides were washed with PBS for 5 min. Nuclei were stained with DAPI (Thermo, 62248) for 15 min at room temperature, followed by a 5-min PBS wash. Slides were mounted with 75 μl of mounting medium and cover slipped for initial DAPI imaging. Coverslips were then removed in PBS, and sections were incubated with primary antibodies against PRG4, CD68, TREM2, and CD31, or with appropriate isotype controls, diluted in the blocking buffer (**Table S5**) for 1 h at room temperature. Sections were washed twice with PBS and incubated with secondary antibodies under the same conditions. Slides were remounted with 75 μl mounting medium for autofluorescence/background imaging. *Dye Inactivation and Multiplexing.* Four signals, including DAPI, were imaged at 20× magnification in regions previously identified during background acquisition. Dye inactivation between staining rounds was performed using 0.1 M NaHCO₃ (pH 11.2, Sigma S6297) and 3% H₂O₂ (Merck, 216763) for three 15-min intervals, with 1-min PBS washes between treatments. After each inactivation step, DAPI was applied to aid computational alignment. This staining and imaging cycle was repeated for seven rounds of primary antibodies. *Multiplexed Immunofluorescent Image Acquisition.* Sections were visualized on the Cell DIVE microscope (Leica/GE) with ImageApp software. A 10× reference scan of each slide was used to select regions of interest. Post-DAPI staining, 20× images were acquired to compensate for autofluorescence in the green (AF488), orange (Cy3), and far-red (Cy5) channels, and to generate virtual H&E images. Primary-secondary staining was used for PRG4 and TREM2, while directly conjugated antibodies were used for CD31 and CD68 (Table S5). Autofluorescent background subtraction was performed after each dye inactivation step. Image alignment was carried out automatically using ImageApp, or manually using anchor cells where significant tissue loss occurred. *Preparation of Final Images* Final image visualization, including isotype controls and primary-stained slides, was performed using QuPath (v0.4.2). As adult and developmental joint sections exhibited different levels of background signal, exposure values for channels containing primary stains are provided in **Table S6.**

### Synovial organoid cells phenotyping and sorting

SOs were digested into single-cell suspensions using Liberase™ Research Grade TM in RPMI (0.15 mg/mL, 0.78 Wünsch units/mL) for 30 min, as described previously (*7, 24*). After incubation, organoids were mechanically dissociated by pipetting up and down at least 10 times using 1000 µL wide-bore tips. Tubes were returned to a shaking incubator for an additional 30 min. The dissociation process was then repeated using narrow-bore tips. After a total of 60 min of digestion, 0.5 mL of fresh digestion buffer was added, followed by further pipetting with narrow-bore tips and a final 30-min incubation. After 90 min, SOs were fully digested into single-cell suspensions with no visible clumps. One ml of cold complete medium (RPMI with 10% foetal calf serum) was added to each 15 mL tube to neutralize the digestion buffer, and cells were centrifuged at 400 × g for 5 min at 4 °C. The supernatant was carefully removed, and cell pellets were gently resuspended in 1 mL of cold FACS buffer (2% FCS, 100 U/mL penicillin/streptomycin, 2 mM Glutamax, 2 mM EDTA) using 1000 µL wide-bore tips to minimize cell damage. Resuspended cells were transferred to 5 mL polypropylene round-bottom FACS tubes in a final volume of 2 mL and centrifuged again. For phenotyping and FACS sorting, we followed previously published protocols (*7, 12*). Briefly, an aliquot of the cell suspension was set aside as an unstained control. Cells were stained with Fixable Viability Dye eFluor 780 (eBioscience) at 1:1,000 in PBS and incubated on ice for 20 min, then washed with FACS buffer and centrifuged at 400 × g for 5 min at 4 °C. The supernatant was removed, and an aliquot was retained for live-dead control. Remaining cells were stained on ice for 30 min with antibodies targeting myeloid, fibroblast, and endothelial populations (all antibodies listed in **Table S3**). SOs lacking immune cells and fluorescence-minus-one (FMO) controls for CD45 were used to define gating values. Cells were washed with 1 mL FACS buffer, resuspended in 250–300 μL, and filtered through a 100-μm cell strainer before analysis or sorting on a FACS ARIAIII (BD Biosciences) using 100- or 130-μm nozzles, respectively.

### Integration of single cell omics of synovial organoids with a reference synovial tissue data set

Synovial organoid cells were labelled with unique sample identifier tags (Sample Tag 1–12) using the BD Human Single-Cell Sample Multiplexing Kit (633781, BD Biosciences), and single cell transcriptomics was performed with BD Rhapsody platform as described above. To integrate scRNAseq data set of organoids with synovial tissue scRNAseq data set, we employed the scANVI framework (Python v3.10.12, Scanpy v1.9.3, scvi-tools v1.0.2), a semi-supervised variant of scVI designed for joint analysis of heterogeneous datasets. CD45⁺ SO scRNAseq data were integrated with synovial tissue myeloid cells from MacDonald et al., 2024 (*12*) while CD45⁻ SO scRNAseq were integrated with FLS from Alivernini et al., 2020 (*7*). Common genes across paired datasets were retained, and the combined data were normalized to median counts per cell, followed by log1p transformation. Highly variable genes (HVGs; 2,500 genes) were selected using the Seurat v3 flavour, accounting for batch effects. The SCVI model was trained with default parameters to learn a shared latent representation. scANVI was then initialized from the SCVI model and fine-tuned using predefined cell-type labels from the respective original studies. The integrated latent space was used for downstream clustering and UMAP visualization. Data were then converted to Seurat for final plots. Clusters markers were identified with FindAllMarkers, (test.use=MAST), as genes with logFC > 0.6, with significant adjusted p-value <0.05 and expressed by greater than 30% of cells in the cluster (‘min.pct’ parameter 0.3).

### Statistical analysis

Detailed statistical methods are provided in each figure legend and in the scRNAseq method sections above.

## List of Supplementary Materials

**Supplementary Materials**

Supplementary Figures: Fig S1 to S8

Supplementary Tables: Tables S1 to S6

## Acknowledgments

We thank Dr Leandro Lemgruber Soares, Mrs Diane Vaughan, and Mrs Alana Hamilton (University of Glasgow) for support in imaging and flow cytometry. We would like to thank Ananya Bhalla (University of Oxford) for her help with CellDIVE. We are grateful to all RA patients, healthy volunteers, and nurse team from the SYNGem Biopsy Unit (Division of Rheumatology and Clinical Immunology, Fondazione Policlinico Universitario A. Gemelli IRCCS – Università Cattolica del Sacro Cuore, Rome) for their contributions.

## Funding

This work was supported by

Versus Arthritis UK grant no. 22072 (M.K-S, I.B.M, C.B)

Versus Arthritis UK grant no. 22253 (M.K-S)

Versus Arthritis UK grant no. 22273 (M.K-S)

Versus Arthritis UK grant no. 23229 (M.K-S, S.A and C.B)

Linea D1, Università Cattolica del Sacro Cuore, no. R4124500654 (S.A.)

Ricerca Finalizzata Ministero della Salute no. GR-2018-12366992 (S.A.)

European Horizon 2020 MSCA-COFUND grant no. 847551 (S.W.T, E.M.L.P.)

Netherlands Organisation for Health Research and Development (ZonMw) Create2Solve grant no. 01142052310007 (S.W.T, M.K-S)

NIH-NIAMS K08AR077037 (K.W)

Burroughs Wellcome Fund Career Awards for Medical Scientists (K.W)

Doris Duke Charitable Foundation Clinical Scientist Development Award (K.W)

---

**Supplementary Figure 1 related to Figure 1.**
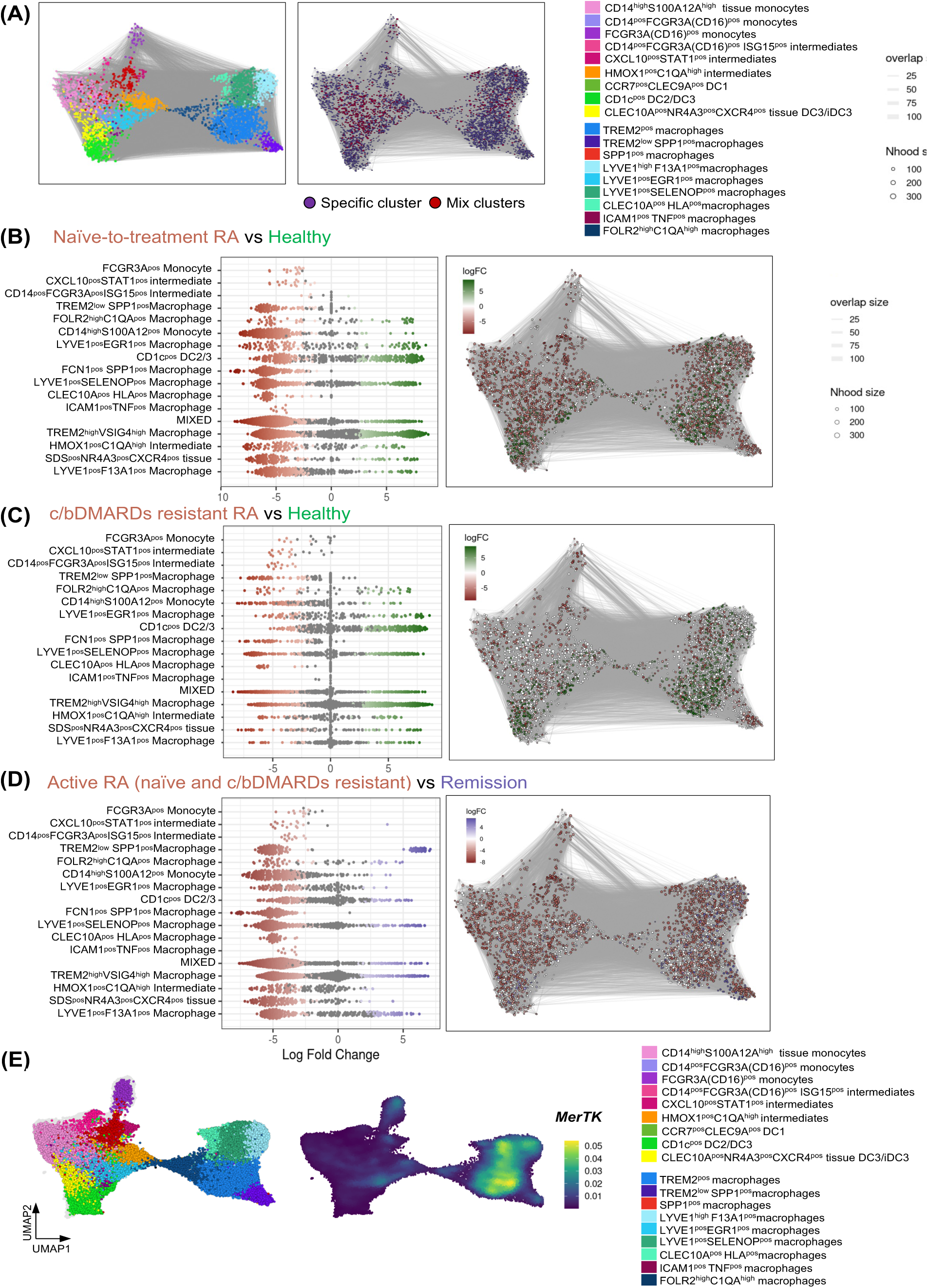
Detailed atlas of macrophage cell clusters in human synovial tissue across different conditions. **(A)** Dataset as in Figure 1 is visualized as MiloR neighbourhood graphs where nodes represent neighbourhoods, coloured by the most represented cell type cluster (left) or by whether >45% of cells within a neighbourhood belong to one specific cluster or not (right). **(B-D)** The changes in synovial tissue myeloid cell composition between healthy controls (n = 11) and naïve-to-treatment RA (n=12) or resistant to c/bDMARDs treatments (n=11) and between active RA (naïve and resistant to treatment) and in RA in sustained remission (n=11) as in Fig.1. Data are visualized as MiloR neighbourhood graphs on the left, where nodes represent neighbourhoods, coloured by their log fold change across conditions. Neighbourhoods with non-differential abundance (FDR > 10%) are coloured white, and node size reflects the number of cells in each neighbourhood. On the right, a beeswarm plot displays the distribution of log fold change across conditions for neighbourhoods containing cells from different cell type clusters. Differential abundance neighbourhoods at FDR ≤ 10% are highlighted in colour. **(E)** Density plots showing MerTK expression in synovial tissue myeloid cell atlas.

**Supplementary Figure 2 related to Figure 1.**
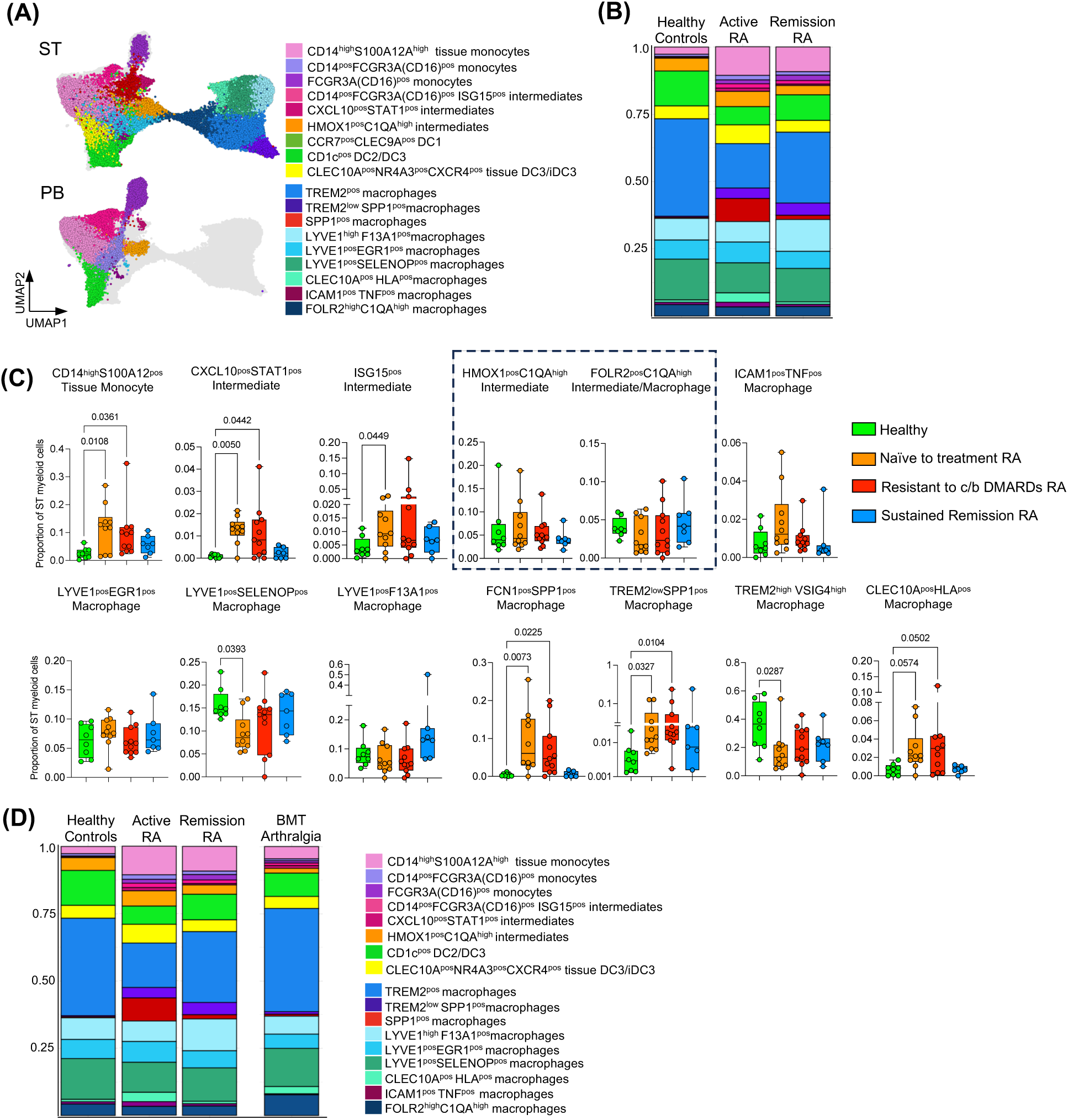
Proportion of distinct synovial tissue myeloid cell clusters between joint conditions. **(A)** UMAP visualization of peripheral blood (PB; n = 5 healthy donors and n = 3 RA patients) and synovial tissue from RA patient samples with active disease, including n = 23 (12 naïve to treatment and 11 resistant to c/bDMARDs) and n = 11 RA in sustained remission (longer than 9 months), as well as from healthy controls (n = 11). Data represent integrated single-cell transcriptomic analysis of synovial tissue (ST) myeloid cells as in Fig.1. **(B)** Stacked plot illustrating the frequency of STM clusters in healthy and RA synovial tissue. **(C)** The proportion of different myeloid cell clusters in synovial tissue differs between healthy controls, active RA, and RA in remission. Data are presented as boxplots showing the median and interquartile range; each dot represents an individual donor/patient. Statistical analysis was performed using the Kruskal–Wallis test with Dunn’s correction for multiple comparisons. Exact p-values are shown on the graphs. The dotted box highlights two related clusters representing an intermediate state between tissue monocytes and macrophages, which in organoids are pooled into a single C1QA^pos^ cluster. **(D)** Stacked plot illustrating the frequency of STM clusters in a bone marrow transplant (BMT) patient in the context of the healthy and RA synovial tissue atlas.

**Supplementary Figure 3 related to Figure 2 and 4.**
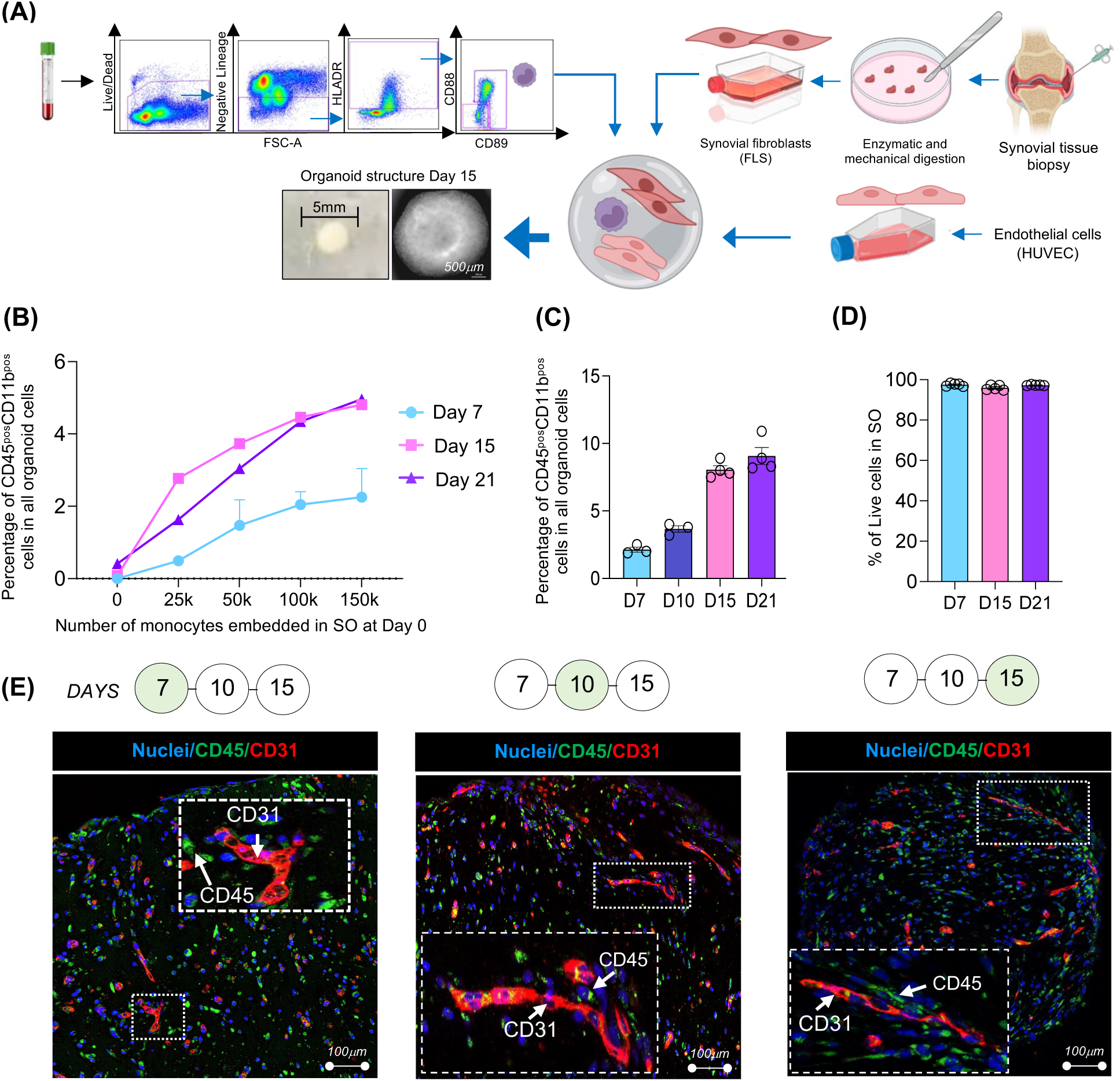
Establishing a human myeloid-stromal synovial organoid system. **(A)** Schematic illustrating the workflow of synovial organoid (SO) generation using synovial fibroblasts, blood monocytes, and endothelial cells in a single droplet of Matrigel as in Fig.2C. **(B)** Percentage of myeloid cell (CD45⁺ CD11b⁺) retrieved from SO with different monocyte numbers embedded at day 0. Data are presented as the mean ± SD (where n = 2 replicates) of the number of myeloid cells across time points (Days 7, 14, 21; total n = 20 SOs from 2 independent experiments). **(C)** Percentage of myeloid cell (CD45⁺ CD11b⁺) retrieved from synovial organoids (SO) seeded with 100K monocytes, showing the number of macrophages at different time points. Bar plots show the mean ± SEM % of myeloid cells within total SO cell number. Data from n = 3–4 synovial organoids across 3 independent experiments. **(D)** Bar plots showing the mean ± SEM of cell viability from n = 6 synovial organoids in 2 independent experiments. **(E)** Representative confocal microscopy images (40x) showing IF staining for CD31 (red), macrophages (CD45, green) and nuclei stained with DAPI (blue) from organoids with FLS derived from biopsies of n=6 patients with active RA at different time points. The insert shows enlarged region of SO. Scale bars, 100 µm.

**Supplementary Figure 4 related to Figure 4.**
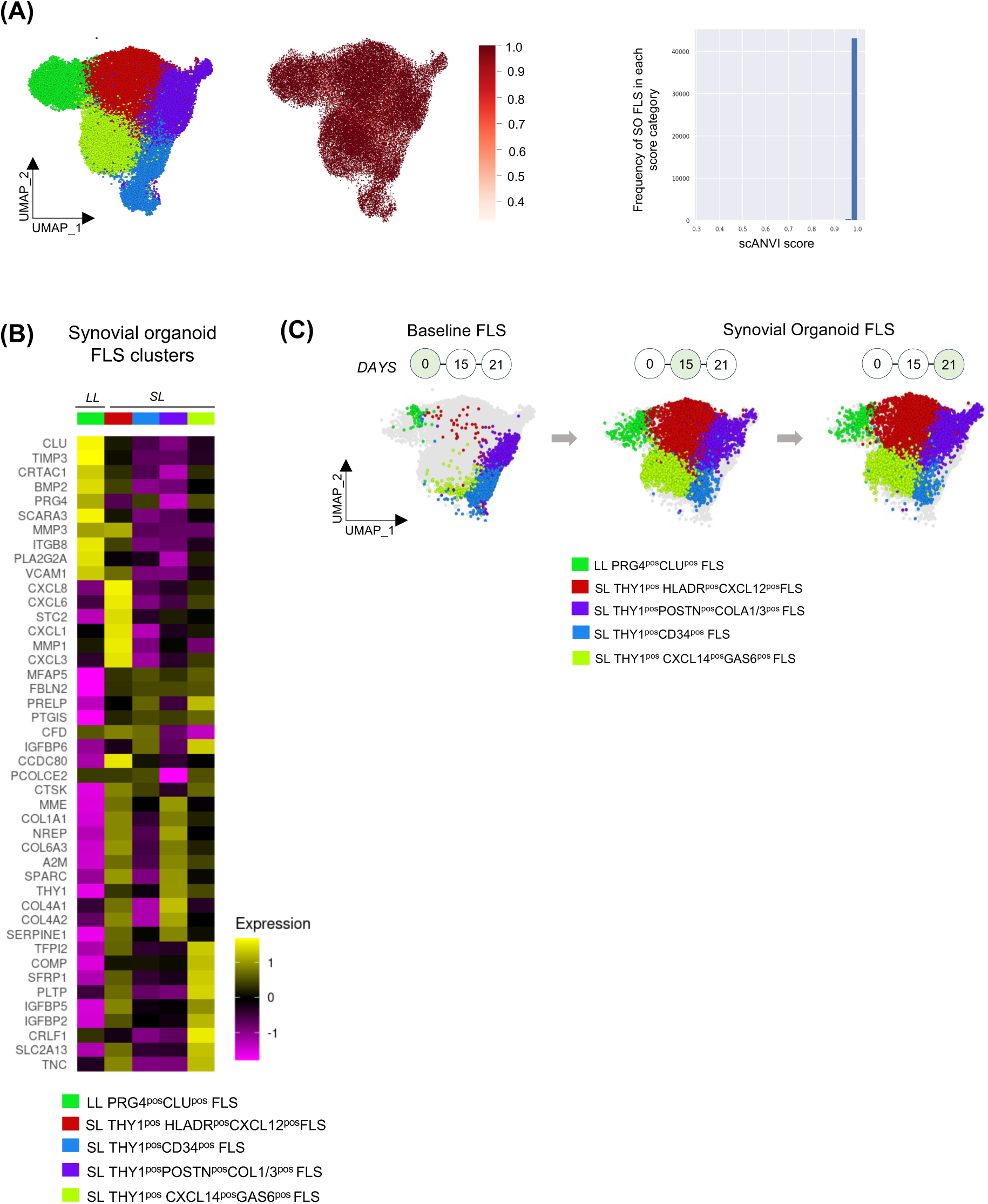
Synovial organoid FLS cluster composition resembles synovial tissue. **(A)** UMAP visualization of integrated single-cell data of synovial organoid (SO) and synovial tissue FLS and respective scANVI prediction score (score_scanvi), as feature plot or frequency plot. The score reflects the model’s confidence (0–1) in cell-type label assignments, with higher values indicating greater confidence in the predicted cell identity of SO cells. (B) Heatmap visualizing the gene expression of the top 10 marker genes of SO FLS clusters. Differentially expressed (DE) genes were determined using MAST and were considered significant if expressed in more than 30% of cells in the appropriate cluster with adjusted p < 0.05 after Bonferroni correction for multiple comparison, average logFC≥0.5. (C) UMAP visualization of baseline fibroblasts and SO fibroblast clusters at day 15 and at day 21 (n = 4 technical replicates) as well as a reference tissue FLS. SO=synovial organoids

**Supplementary Figure 5 related to Figure 4.**
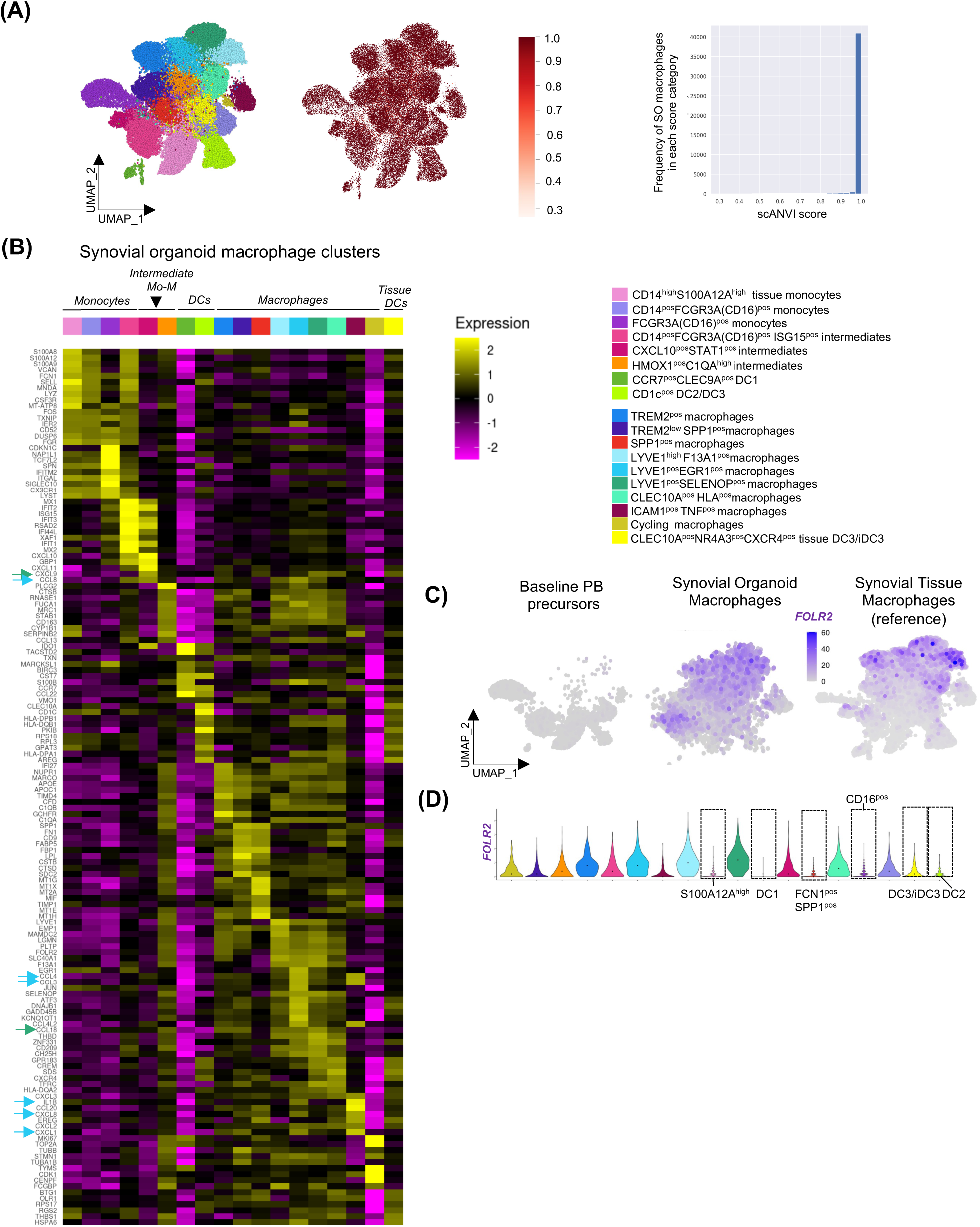
Synovial organoid macrophages (SOM) resemble synovial tissue macrophage (STM). **(A)** UMAP visualization of integrated single-cell data of synovial organoid (SO) and synovial tissue myeloid cells and respective scANVI prediction score (score_scanvi), as feature plot or frequency plot. The score reflects the model’s confidence (0–1) in cell-type label assignments, with higher values indicating greater confidence in the predicted cell identity of SO cells. **(B)** Heatmap visualizing the gene expression of the top 10 marker genes of SO myeloid cells clusters. Differentially expressed (DE) genes were determined using MAST and were considered significant if expressed in more than 30% of cells in the appropriate cluster with adjusted p < 0.05 after Bonferroni correction for multiple comparison, average logFC≥0.5. **(C)** UMAP feature plot showing the expression levels of the tissue macrophage marker FOLR2 in blood precursors, synovial organoids and in reference synovial tissue macrophages. **(D)** Violin plots showing FOLR2 expression in synovial organoid myeloid cell clusters. Dotted frames indicate clusters negative for FOLR2 expression. Arrows indicate gene of interest in perivascular LYVE1^pos^ clusters. SO=synovial organoids.

**Supplementary Figure 6 related to Figure 4.**
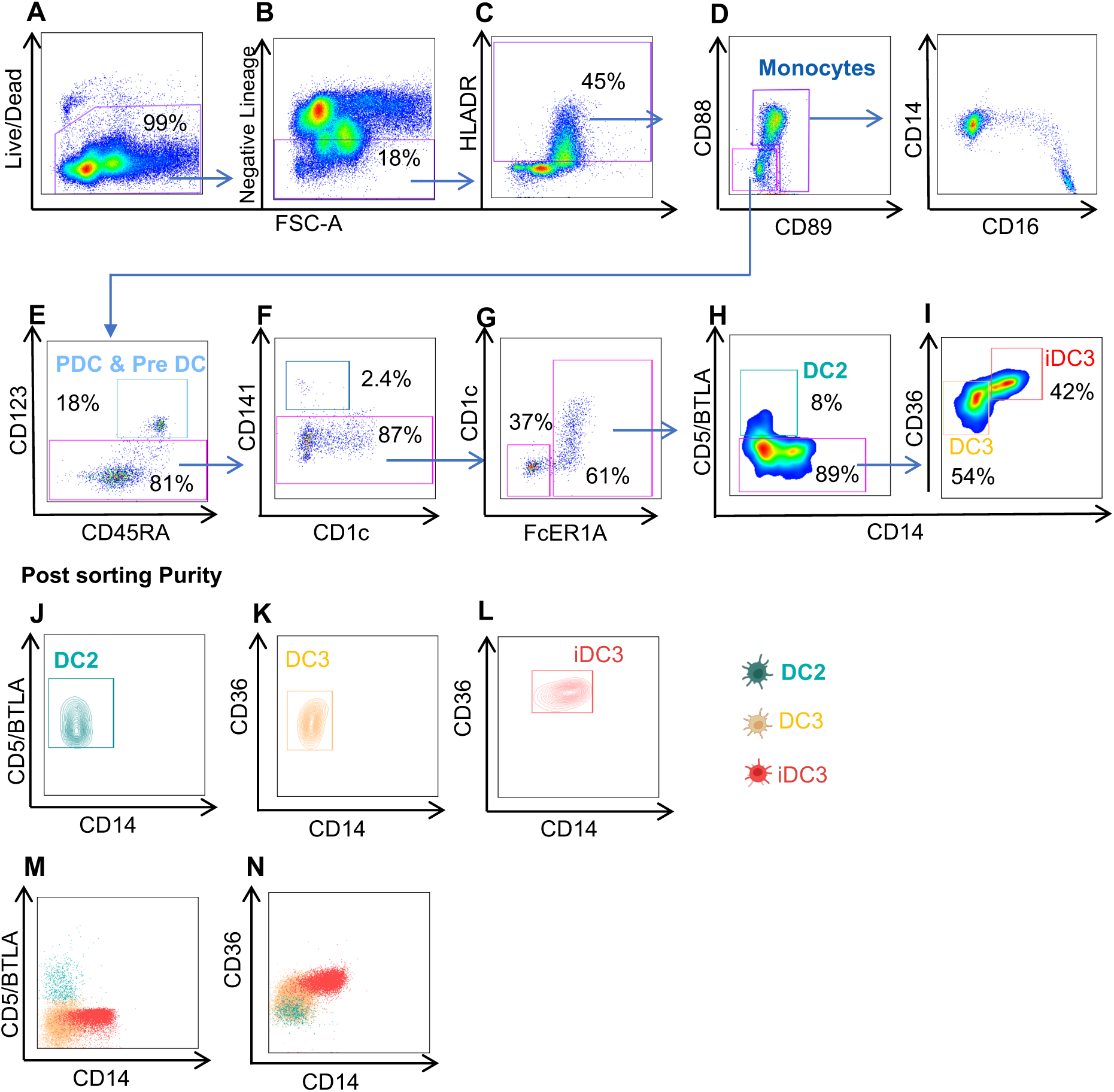
Representative gating strategy for the sorting of healthy PB myeloid populations (monocytes, DC2, DC3 and iDC3) prior to SNP-based organoid generation. Live cells were gated (A), lineage-positive CD3, CD19, CD15, CD117, CD56 cells were excluded (B), and HLA-DR-positive cells were gated (C). Monocyte were gated based on CD88 and CD89 expression (D). After exclusion of PDC, Pre-DC, and DC1 (E & F), myeloid DCs were gated based on CD1c and FcER1A expression (G). DC2 cells were sorted based on CD5/BTLA (H), while DC3 and iDC3 based on distinct expression of CD14 and CD36 (I). J-N validate the post-sorting purity. All the gates were based on unstained cells or Fluorescence Minus One (FMO) control.

**Supplementary Figure 7 related to Figure 4.**
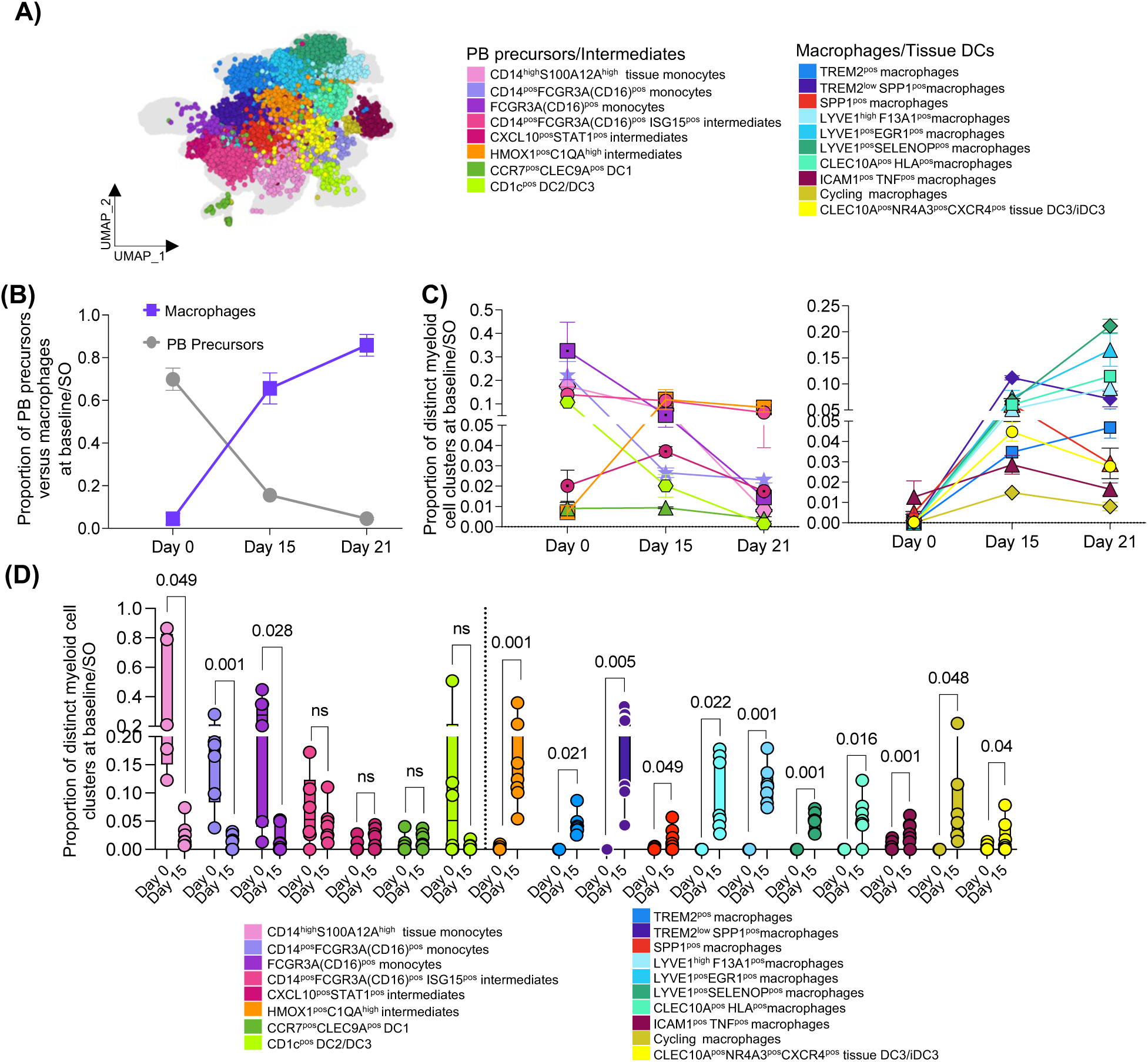
Frequency of myeloid cell precursors and distinct myeloid tissue cell clusters derived from them in human synovial organoids (SO). **(A)** Overview of integrated SO myeloid clusters as in Fig.4D. **(B)** Proportional distribution of PB precursors versus macrophage clusters in SO from data as in (Fig.4J) over time. Cells that mapped with blood cells were defined as PB monocytes/DC and cells that uniquely mapped to synovial tissue were categorized as tissue myeloid cells. Data are presented as a connected Mean ± SD of 2 (baseline) and 4 (SO) technical replicates. **(C)** Kinetics of the development of different synovial tissue macrophage clusters from blood monocytes (Day 0) in synovial organoids. Integrated (scANVI) scRNAseq of the myeloid compartment of synovial organoids at days 15 and 21 and matched baseline monocytes (Day 0). Data are presented as mean ± SEM of n = 4 technical replicates. **(D)** Proportional distribution of monocyte, dendritic cells and macrophage clusters at baseline (Day 0) and in synovial organoids (SO) at day 15, based on n = 7 experiments as in Figure 4 D-E. Data are presented as the median with the interquartile range. Each dot represents a patient-derived SO. A paired T-test between baseline monocytes (Day 0) and matched myeloid cells at Day 15 organoids, with exact p-values displayed on the graph.

**Supplementary Figure 8 related to Figure 6.**
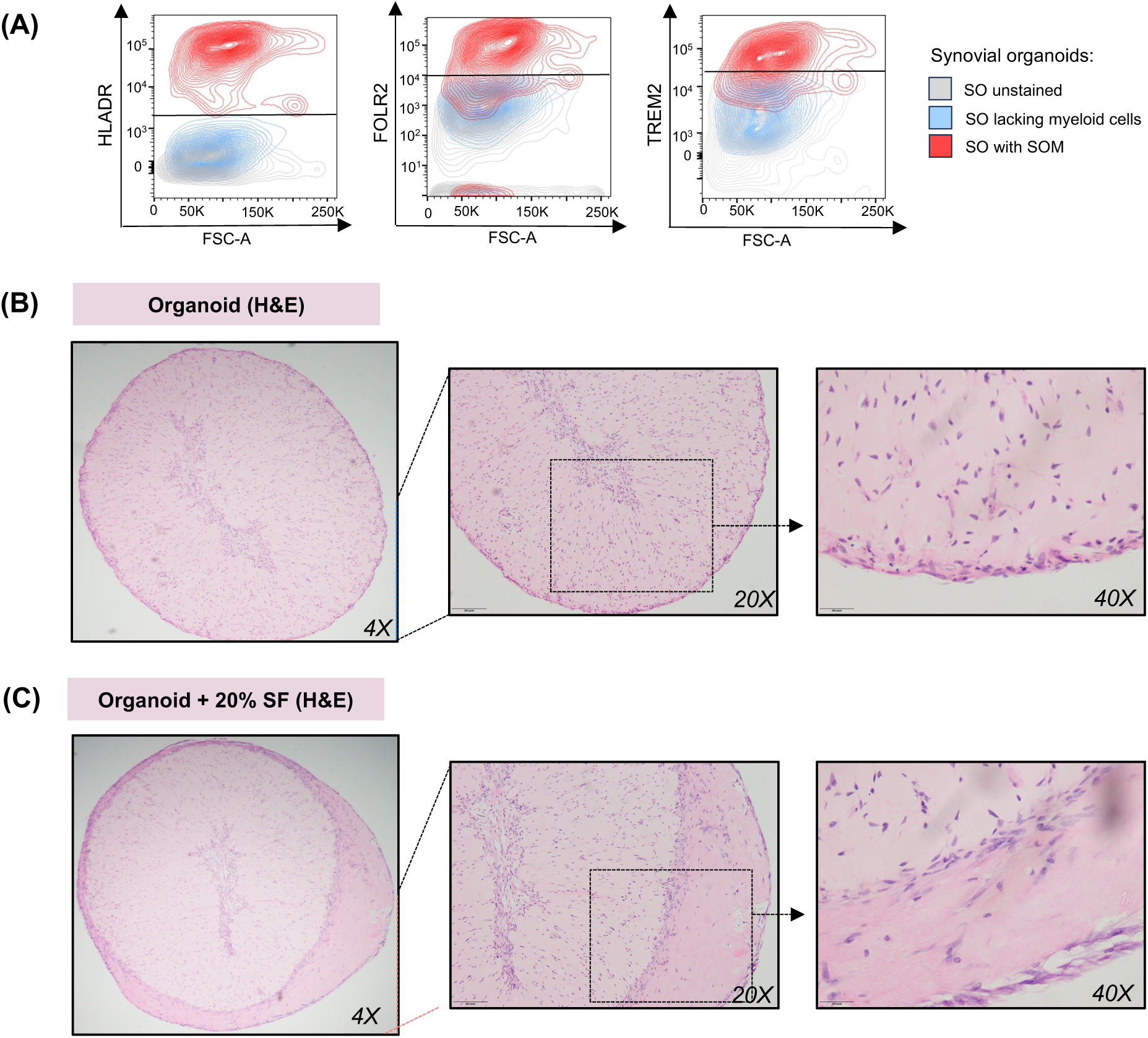
Synovial organoid topography and gating strategy for synovial organoid macrophages. **(A)** Representative flow cytometry contour plots showing expression of HLADR, FOLR2 and TREM2 by synovial organoid macrophages. SO were digested at day 15. Unstained SO, grey, SO lacking myeloid cells, blue and SO stained with selected STM markers in red. Data are generated from FLS derived from n=5 RA patients’ biopsies with active diseases in five independent experiment with 2-4 technical replicate. **(B)** (**C)**The topography of synovial organoids stimulated with synovial fluid (SF). Representative H&E staining of synovial organoid (SO) structure in the absence (B) or in the presence of 20% synovial fluid (SF) for last 7 days of 15 days organoid cultures. Data were generated in 3 independent experiments. SF=pooled cell-free synovial fluid from n=5 patients.

## Author contributions

**A.E.:** (i) contributed to the study concept and manuscript writing, (ii) established the myeloid organoid system **(Fig. 2C),** (iii) established and analysed live imaging of organoids **(Fig. 2G),** (iv) contributed to the developing the gating strategy for PB monocyte and DC cluster sorting **(Fig. S6)** for organoid inclusion, (v) performed digestion, phenotyping, sorting, and analysis of synovial organoid cells and validated synovial organoid spatial topography **(Figs. 2D–F, 3, 4, 5, 6; Figs. S3, S8),** and (vi) performed staining and analysis of human embryonic joints with J.F. **(Figs. 2A–B).**

**D.S.:** (i) contributed to the study concept and manuscript writing, (ii) developed the gating strategy for PB monocyte and DC cluster sorting for organoid inclusion **(Fig. S6),** (iii) deconvoluted condition-specific myeloid cell cluster abundance in synovial tissue datasets **(Fig. 1D; Fig. S1),** (iv) contributed to organoid live imaging analysis **(Fig. 2G),** (v) performed sorting and phenotyping of synovial organoid cells, conducted and analysed scRNAseq data of synovial organoids, integrated them with synovial tissue data, and established the SNP-based cell tracking analysis system **(Figs. 4, 5A–G; Figs. S3A–D, S4, S5, S7, S8A).**

**L.M.:** (i) contributed to the study concept and manuscript writing, (ii) analysed synovial tissue myeloid cell datasets, established the synovial tissue myeloid cell nomenclature, and analysed and integrated with synovial tissue, the scRNAseq data from patients who underwent bone marrow transplantation **(Figs. 1A–C, 1E, 1G–H; Fig. S2),** (iii) performed synovial organoid experiments with A.E., and synovial organoid scRNAseq and analysis **(Figs. 4C–E),** (iv) assisted with analysis of multiple myeloid precursors organoid scRNAseq datasets **(Fig.4G-I)**, and (v) contributed to the development the gating strategy for PB monocyte and DC cluster sorting **(Fig. S6).**

**J.F.:** (i) performed staining and analysis of human embryonic joints with A.E. **(Figs. 2A–B)** and

1. (ii) contributed to the concept of the SNP-based cell fate organoid system **(Figs. 4F–I),** and
2. (iii) contributed to the development the gating strategy for PB monocyte and DC cluster sorting **(Fig. S6).**

**Z.K.:** (i) established the scANVI pipeline for integration of organoid and synovial tissue scRNAseq datasets **(Figs. 4C–D; Fig. S4).**

**L.A.C.: (**i) contributed to flow cytometry validation of organoid scRNAseq data **(Fig. 5B)** and (ii) collected clinical information of the RA patients, and (iii) performed RNAscope on human synovial tissue biopsies **(Fig.1F)**

**T.S.:** performed scRNAseq of the SNP-based cell fate organoid system **(Fig. 4G). M.D.:** assisted with synovial organoid culture.

**E.P. and S.W.T.:** (i) provided synovial fluid from five RA patients and (ii) contributed the concept of incorporating SF into the organoid system **(Fig. 6).**

**S.B.:** analysed live imaging data of organoids **(Fig. 2G).**

**K.W.:** shared the FLS/EC organoid protocol, which was modified in this study for use with myeloid precursors.

**S.G.D. and C.B.:** (i) contributed to the concept synovial niche and (ii) provided human embryonic joints and expertise on the embryonic joint topography **(Fig.2A-B)**

**S.P. M.C.C. and S.D.:** (i) provided human embryonic joints and expertise on the embryonic joint topography (Fig.2A-B)

**D.W.:** contributed to staining and analysis of human embryonic joints **(Fig.2A-B).**

**C.D.M. and D.C.:** (i) performed synovial tissue collections, (ii) tissue sample handling, (iii) generated FLS cultures, (iv), performed Immunofluorescent staining **(Fig.3A),** and (v) RNAscope on synovial tissue biopsies, together with L.A.C **(Fig. 1F).**

**D.B.:** contributed to clinical assessment of patients and synovial tissue collection (Fig.1-6)

**M.G, L.P., V.P.:** performed clinical assessment for condition annotations in active and remission cohorts **(Fig.1-6)**

**Ch.M.:** contributed to manuscript writing.

**I.B.M. and D.T:** assisted with data interpretation.

**M.A.D.A.:** provided clinical setting.

**S.A. and M.K.S.:** initiated the study concept, designed and supervised the overall work, and wrote the manuscript.

## Data and materials availability

Single cell data sets will be available upon manuscript acceptance from ArrayExpress.

All single cell data transcriptomics codes, and protocols used in this study are available upon request to corresponding author: Mariola Kurowska-Stolarska

## Supplementary Tables

**Table S1:** Demographic, clinical and immunological characteristics of Rheumatoid arthritis patients whose synovial tissue biopsies were used to generate synovial organoids.

|  |  |
| --- | --- |
|  | **Active RA**<br>n=10 |
| **Female sex, n(%)** | 8 (80) |
| **Age, years (mean ± SEM)** | 65.3 ± 3.1 |
| **Disease Activity Score, (mean ± SEM)** | 5.7 ± 0.4 |
| **ACPA positivity, n(%)** | 10 (100%) |
| **RF positivity, n(%)** | 10 (100%) |
| **Concomitant treatments:** | Naive, n=4<br>c/b-DMARDs, n=6 |

**ACPA:** Anti-Citrullinated Peptide Antibodies; **RF:** Rheumatoid Factor.
**c/b-DMARDs:** conventional/biologic-Disease Modifying Anti-Rheumatic Drugs.

**Table S2:** Demographic, clinical and immunological characteristics of Rheumatoid arthritis patients whose synovial biopsies were used to extend synovial tissue myeloid cell atlas

|  | **Active RA**<br>n=4 | **Sustained**<br>**Remission RA**<br>n=5 | **Healthy**<br>n=7 |
| --- | --- | --- | --- |
| **Female sex, n(%)** | 3 (75%) | 3 (60%) | 0 (0%) |
| **Age, years (mean ± SEM)** | 55 ± 8.4 | 44 ± 2.1 | 37 ± 5.8 |
| **Disease Activity Score, (mean ± SEM)** | 5.3 ± 0.4 | 1.6 ± 0.2 | N/A |
| **ACPA positivity, n(%)** | 3 (75%) | 4 (80%) | N/A |
| **RF positivity, n(%)** | 2 (50%) | 4 (80%) | N/A |
| **Concomitant treatments:** | Naïve, n=2<br>c/b-DMARDs, n=2 | c/b-DMARDs,<br>n=5 | N/A |

**ACPA:** Anti-Citrullinated Peptide Antibodies; **RF:** Rheumatoid Factor.
**c/b-DMARDs:** conventional/biologic-Disease Modifying Anti-Rheumatic Drugs.

**Table S3.** Details of the antibodies used for sorting and phenotyping peripheral blood precursors and synovial organoid cells.

| **Antibodies for sorting peripheral blood monocytes prior to organoid generation** |  | **Cat No./ Source** |
| --- | --- | --- |
| **1** | APC anti-human **HLA-DR** | 307610 /Biolegend |
| **2** | Brilliant Violet 650 anti-human **CD14** | 301830/Biolegend |
| **3** | Brilliant Violet 510 anti-human **CD16** | 360730/Biolegend |
| **4** | PE/CY7 anti-human **CD88** | 344308/Biolegend |
| **5** | AF700 anti-human **CD89** | 354118/Biolegend |
| **6a** | FITC anti-human **CD15** (SSEA-1) dump channel | 323004/Biolegend |
| **6b** | FITC anti-human **CD19**-dump channel | 302206/Biolegend |
| **6c** | FITC anti-human **CD20**-dump channel | 302304/Biolegend |
| **6d** | FITC anti-human **CD117** (c-kit)-dump channel | 313232/Biolegend |
| **6e** | FITC anti-human **CD56** (NCAM)-dump channel | 304604/Biolegend |
| **6f** | FITC anti-human **CD3**-dump channel | 300440/Biolegend |
| **Antibodies used for sorting/phenotyping fibroblasts and myeloid cells from synovial organoids** |  | **Cat No./ Source** |
| **1** | Brilliant Violet 711 anti-human **CD45** | 304050/Biolegend |
| **2** | Brilliant Violet 785 anti-human **HLA-DR** | 307642/Biolegend |
| **3** | Brilliant Violet 510 anti-human **CD64** | 305028/Biolegend |
| **4** | AF700 anti-human **CD11b** | 301356/Biolegend |
| **5** | APC anti-human **Folate Receptor b** | 391706/Biolegend |
| **6** | PE anti-human **TREM2** | FAB17291P/R&D Systems |
| **7** | APC anti-human **SPP1** | 50-9096-42/Invitrogen |
| **8** | PE/CY7 anti-human **MERTK** | 367604/Biolegend |
| **9** | PE/Dazzle594 anti-human **CD48** | 562717/BD Biosciences |
| **10** | PE/CY7 anti-human **CD90** | 328124/Biolegend |
| **11** | PE/Dazzle594 anti-human **PDPN** | 337028/Biolegend |
| **12** | AF700 anti-human **CD31** | 303104/Biolegend |

**Table S4.** Details of antibodies used for sorting DC subsets and monocytes from peripheral blood of healthy donors for SNP-based synovial organoid system.

|  | Antibody | Cat No./ Source |
| --- | --- | --- |
| **1** | Brilliant Violet 421 anti-human **CD5**/Brilliant Violet 421 anti-human **BTLA** | 300626/Biolegend<br>344512/Biolegend |
| **2** | Brilliant Violet 510 anti-human **CD45RA** | 304142/Biolegend |
| **3** | Brilliant Violet 605 anti-human **FcER1A** | 334628/Biolegend |
| **4** | Brilliant Violet 650 anti-human **CD14** | 301836/Biolegend |
| **5** | Brilliant Violet 711 anti-human **CD163** | 333630/Biolegend |
| **6** | Brilliant Violet 785 anti-human **HLA-DR** | 307642/Biolegend |
| **7** | PE anti human **CD141** | 559781/BD |
| **8** | PE/CY5 anti-human **CD16** | 302010/Biolegend |
| **9** | PE/CY7 anti-human **CD88** | 344308/Biolegend |
| **10** | PE/Dazzle 594 anti-human **CD123** | 306034/Biolegend |
| **11** | APC anti-human **CD1c** | 331524/Biolegend |
| **12** | AF700 anti-human **CD89** | 354118/Biolegend |
| **13** | PerCP_Cy5.5 _anti-human **CD36** | 336224/Biolegend |
| **14a** | FITC anti-human **CD15** (SSEA-1) dump channel | 323004/Biolegend |
| **14b** | FITC antihuman **CD19** dump channel | 302206/Biolegend |
| **14c** | FITC anti-human **CD117** (c-kit) dump channel | 313232/Biolegend |
| **14d** | FITC antihuman **CD3** dump channel | 300440/Biolegend |
| **14e** | FITC anti-human **CD56** (NCAM) dump channel | 304604/Biolegend |

**Table S5.** Primary and secondary antibodies used to map the fibroblast and immune cells in human synovial tissue biopsies, embryonic joints and synovial organoids.

| Primary Antibodies | Working Dilution / Concentration | Cat No./ Source | Secondary antibodies |
| --- | --- | --- | --- |
| Rabbit anti-human **COL1A** | 1:100 | HPA011795/Sigma | MP-7401/ ImmPRSS HRP Horse Anti Rabbit |
| Rabbit anti-human **COL3A** | 1:100 | HPA007583/Sigma | MP-7401/ ImmPRSS HRP Horse Anti Rabbit |
| Rabbit anti-human **PDPN** | 1:200 | HPA007534/Sigma | A-21245/ Goat anti-Rabbit IgG Alexa Fluor 647 (1:100) |
| Rabbit anti-human **MMP3** | 1:50 | ab52915 /Abcam plc | A-21245/ Goat anti-Rabbit IgG Alexa Fluor 647 (1:100) |
| Rabbit anti-human **CD31** | 1:100 | Ab76533 /Abcam plc | A-21245/ Goat anti-Rabbit IgG Alexa Fluor 647 (1:100) |
| Mouse anti-human **CD31** | 1:200 | Abab9498 /Abcam plc | A-212236/ Goat anti-Mouse IgG Alexa Fluor 647(1:100) |
| Rabbit anti-human **LYVE1** | 1:200 | HPA042953 / Sigma | A11008/ Goat anti-Rabbit IgG Alexa Fluor 448 (1:100) |
| Rabbit anti-human **SPP1** | 1:100 | ab101492/Abcam plc | A-21245/ Goat anti-Rabbit IgG Alexa Fluor 647 (1:100) |
| Rabbit anti-human **CD90** | 1:100 | Ab226123/Abcam plc | A-21245/ Goat anti-Rabbit IgG Alexa Fluor 647 (1:100) |
| Goat anti-human **TREM2**<br>(Used in IF / Cell DIVE) | 1:50 | ab85851 /Abcam plc | A-21447/ Donkey anti-Goat IgG Alexa Fluor 647 (1:100) |
| Mouse anti-human **PRG4**<br>(Used in IF / Cell DIVE) | 1:100 | MABT401/ Sigma | A-212236/ Goat anti-Mouse IgG Alexa Fluor 647(1:100) (**IF**)<br>A-21202/ Donkey anti-mouse IgG Alexa Fluor 448 (1:100) (**Cell DIVE**) |
| Mouse anti-human **CD45** | 1:200 | NBP2-44863/ Novus-Bio | A-21202/ Donkey anti-mouse IgG Alexa Fluor 448 (1:100) |
| Mouse anti-human **CD68** | 1:40 | M087629-2/ Dako, Cambridgeshire, UK | A-21202/ Donkey anti-mouse IgG Alexa Fluor 448 (1:100) |
| Rabbit anti-human **CD68**<br>(Used in Cell DIVE) | 10 µg/mL | ab280860 / Abcam plc | Directly conjugated to AF555 |
| Mouse anti-human **CD31**<br>(Used in Cell DIVE) | 10 µg/mL | NBP2-33154AF647 / Novus-Bio | Directly conjugated to AF647 |

**Table S6.**
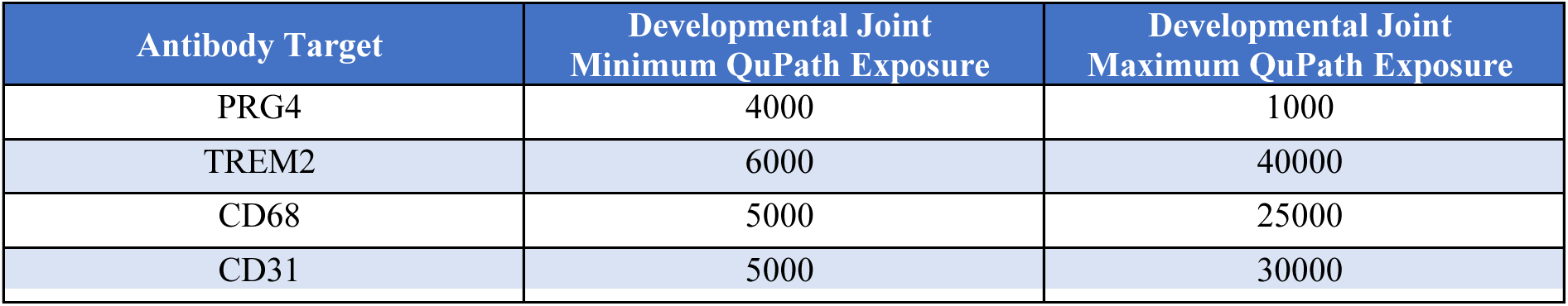
Minimum and maximum exposure values used in QuPath analysis software for images displayed in Figure 2A-B.

